# A peptide strategy for inhibiting different protein aggregation pathways in disease

**DOI:** 10.1101/2022.10.22.513060

**Authors:** Tommaso Garfagnini, Luca Ferrari, Margreet B. Koopman, Sem Halters, Eline Van Kappel, Guy Mayer, Madelon M. Maurice, Stefan G. D. Rüdiger, Assaf Friedler

## Abstract

Protein aggregation correlates with many human diseases. Protein aggregates differ in shape, ranging from amorphous aggregates to amyloid fibrils. Possibly for such heterogeneity, strategies to develop effective aggregation inhibitors that reach the clinic failed so far. Here, we present a new strategy by which we developed a family of peptides targeting early aggregation stages for both amorphous and fibrillar aggregates of proteins unrelated in sequence and structure. Thus, they act on dynamic precursors before a mechanistic differentiation takes place. Using a peptide array approach, we first identified peptides inhibiting the predominantly amorphous aggregation of a molten globular, aggregation-prone protein, a thermolabile mutant of the Axin tumor suppressor. A series of optimization steps revealed that the peptides activity did not depend on their sequences but rather on their molecular determinants. The key properties that made a peptide active were a composition of 20-30% flexible, 30-40% aliphatic and 20-30% aromatic residues, a hydrophobicity/hydrophilicity ratio close to 1 and an even distribution of residues of different nature throughout the sequence. Remarkably, the optimized peptides also suppressed fibrillation of Tau, a disordered protein that forms amyloids in Alzheimer’s disease, and entirely unrelated to Axin. Our compounds thus target early aggregation stages, independent of the aggregation mechanism, inhibiting both amorphous and amyloid aggregation. Such cross-mechanistic, multi-targeting aggregation inhibitors may be attractive lead compounds against multiple protein aggregation diseases.

## Introduction

The aberrant aggregation of proteins leads to the formation of soluble or insoluble agglomerations of no longer functional protein^1^. Any protein can aggregate in response to molecular stresses that disrupt the proteostatic equilibrium^2^. Mutations, post-translational modifications, thermal or oxidative shocks destabilize protein structure, modify the exposed surface and establish aberrant protein-protein interactions (PPIs) capable of triggering aggregative events^2,3^. Such precursors of protein aggregation constitute a dynamic pool of transient, metastable states and act as initiators in the early stages of aggregation of all kinds^1^ (Fig. 1). Aggregation then proceeds via different mechanisms through numerous types of heterogeneous intermediate species and results in aggregates with distinct properties: amorphous aggregates lack structural order and defined shape; native-like aggregates retain most of protein native folding while displaying varied morphologies; amyloid aggregates are unbranched fibrils formed by proteins extensively enriched in β-sheet and assembled into a highly packed cross-β superstructure^1^.

**Figure 1.**
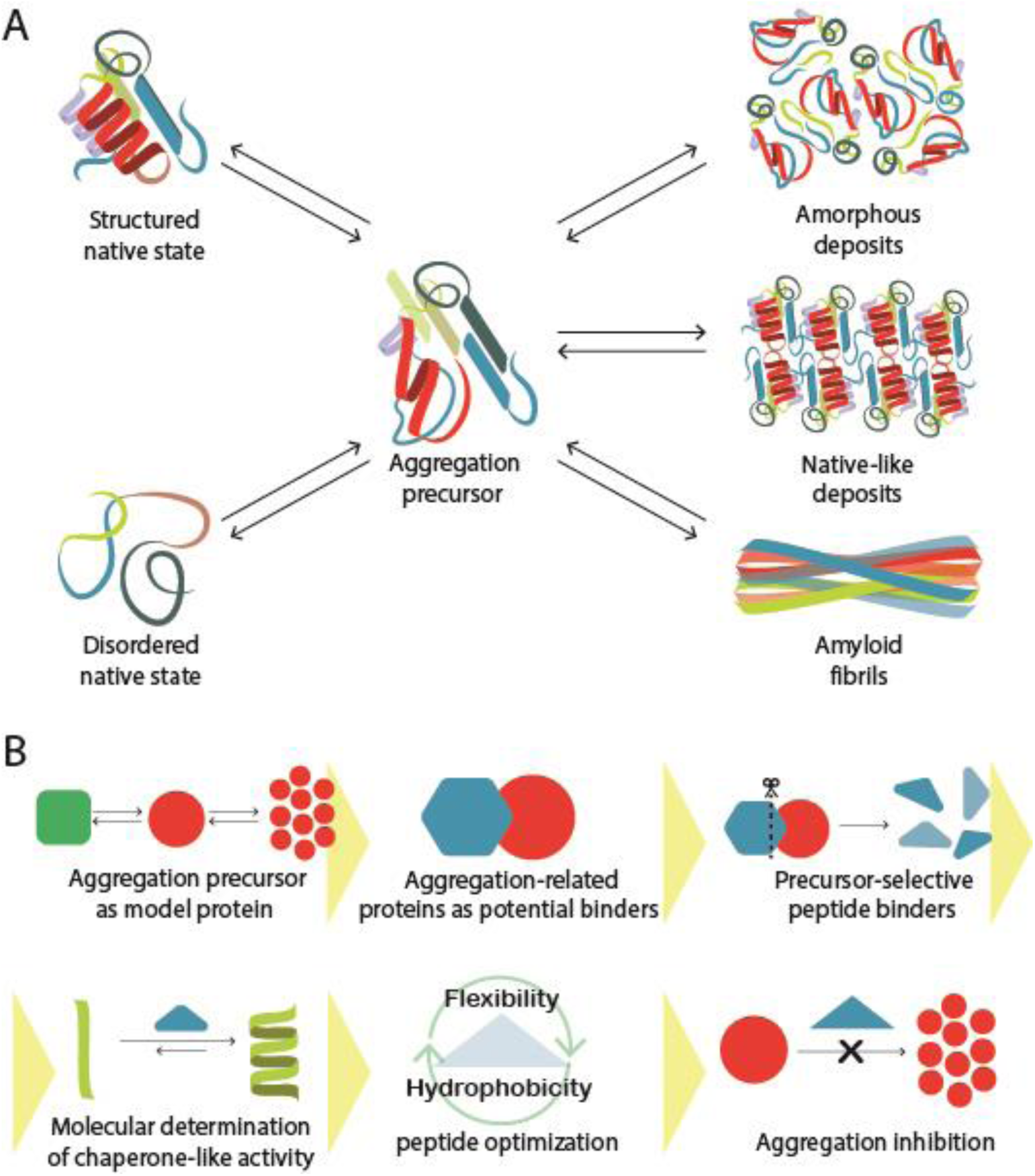
A scheme of protein aggregation and of the targeting strategy presented in the current study. A) Structured and disordered proteins access aggregation precursor states that trigger a process of aberrant assembly. The mechanistic differentiation after early aggregative events leads to amorphous aggregates, native-like aggregates or amyloid aggregates; B) Our strategy for general targeting of aggregation precursor states (red) employed selected peptides derived from precursor-interacting proteins (cyan). Their sequences were optimized via a series of rational design steps leading to peptides that interact with the target proteins in a chaperone-like manner.

The appearance of protein aggregates has various pathological outcomes, such as cytotoxicity, immunogenic responses and cell homeostasis disruption by inactivating proteins that orchestrate key regulatory processes^2,4,5^. This results in various diseases ranging from some cancer phenotypes, to neurological disorders such as Alzheimer’s, Parkinson’s and Huntington’s diseases and amyotrophic lateral sclerosis, or systemic diseases like type II diabetes, liver, cardiac, kidney amyloidosis and cataract^1–4,6^. Therefore, developing tools for studying and inhibiting protein aggregation in disease is of extreme importance.

Many peptide aggregation inhibitors were developed for specifically targeting one sequence identified as the main aggregation hotspot of the target protein^7–15^. Multi-targeting is sometimes purposely pursued to achieve specificity towards two proteins at a time, based on sequence/structure homology of the hotspots or the existence of cross-seeding interfaces^16–18^. Yet, aggregation may involve multiple hotspots or occur through different pathways, each one potentially correlating with distinct pathological outcomes^19^. So far, despite an extensive research focus in the past decades, the strategy of addressing protein aggregation in disease did not make a clinical impact yet. Only a few Aβ-targeting antibodies reached or passed stage 3 of Alzheimer’s disease clinical trials, proving that the targeting of aggregation can be used for therapy^20^. However, being very expensive and not orally bioavailable, antibodies are not ideal drugs. No small molecule or peptide inhibitor of protein aggregation had reached the clinic, and only two small molecule candidates (anle138b and UCB0599) decreasing α-synuclein aggregation are at phase 1 of Parkinson’s disease clinical trials^21–25^. A possible reason can be that the peptide inhibitors developed so far are highly specific for certain amyloidogenic sequences, and do not address other pro-aggregating hotspots nor other pathologically relevant aggregates^5,6^. Moreover, such inhibitors have been even more optimized for binding one hotspot, so they cannot be further developed into active lead compounds. Specificity is hence not necessarily an advantage but is rather an intrinsic limitation that impairs the ability of current inhibitors to antagonize aggregation efficiently.

Interestingly, in the cell the molecular chaperones use a similar principle that overcomes too much selectivity. Chaperones are involved in both protein folding and protein disaggregation. It is the conserved Hsp70 chaperone family that is part of the general folding pathway of the cell and disaggregates both amorphous and fibrillar proteins^26–30^. This means that the same chaperone machine can recognize both amorphous and fibrillar aggregates, and possibly their precursors. Such proteins do not have any sequence similarity. Thus, we assumed it may be an attractive strategy to mimic the physiological situation and develop compounds with broad specificity, acting on both amorphous and fibrillar aggregation.

Here we describe a new approach to overcome the inherent limitations described above by rationally designing peptide aggregation inhibitors of both amorphous and amyloid aggregation (Fig. 1). To achieve this aim, we implemented a multi-protein recognition and targeting strategy inspired by the same recognition principles of molecular chaperones. Our design strategy was based on the search for the molecular signature of anti-aggregation activity encrypted in the amino acid sequence, rather than selecting one hotspot as a mold for shaping specific inhibitors. We used the cancer-related L106R mutant in the RGS domain of the Axin protein (RGS L106R), as a model for a target protein that forms predominantly amorphous, soluble nanoaggregates upon destabilization^31,32^. We screened RGS L106R for binding a library of aggregation-related peptides, and the sequences of the peptides that inhibited aggregation were analyzed to find common traits. We discovered that the inhibitory activity is inherently encoded in the molecular determinants of the peptide sequence without involving consensus patterns. The determinants were optimized through a series of rational design steps that resulted in peptides with enhanced activity, proving that the sequence is a starting point for improvement and optimization, adaptable to various protein clients. As a proof of concept, the peptides also inhibited aggregation of Tau, an intrinsically disordered protein that undergoes amyloid aggregation in a radically different mechanism^33,34^. Such determinants hence provide a novel principle for the rational design of peptides inhibitors that potentially oppose the aggregation of any protein, irrespective of sequence, structure or aggregation mechanism, in a similar manner as chaperones do. Our work paves the way to rendering chaperone-like activity at the peptide scale for drug discovery against any protein aggregation disease.

## Results

### A general strategy for designing aggregation inhibitors

We set out to develop an inhibitory strategy that targets the aggregation precursor state. We hypothesized that since the aggregation precursors possess similar biophysical and chemical properties potentially shared by all mechanisms, they could be collectively targeted by designed peptides with specific chemical properties that enable binding and inhibition. Our strategy (Fig. 1B) was thus first to develop inhibitors of amorphous aggregation, optimize their molecular properties and then test their ability to inhibit amyloid aggregation, which follows a completely different mechanism.

We isolated the early aggregative interactions from those that are mechanism-specific by using Axin RGS L106R as a model protein, a molten globule that natively resides in an aggregation precursor state and forms soluble, predominantly amorphous nanoaggregates of 4-5 molecules without structural rearrangements^31,32^. Within the RGS sequence, the residues ^116^DFWFACTGF^124^ display the highest intrinsic propensity to aggregate, and experimental evidence confirms that this stretch leads RGS L106R aggregation *in vitro* and *in vivo*^31^. We then tailored a peptide targeting strategy to shield the aberrant interactions of such an aggregation hotspot and prevent RGS L106R aggregation. To find peptides that are selective towards RGS L106R, we designed an array of peptides derived from a pool of proteins with potential RGS L106R binding sequences: ***i*)** Axin, that may participate in RGS aggregation through other parts of its sequence^31^; ***ii*)** amyloid-forming/binding proteins, which may bind via cross-recognition between aggregation-prone strands, such as Aβ_42_, HypF-N, SsoAcP, mAcP, the cysteine rich domains (CRDs) of Frizzled 1/3/4/5/6/9 and DKK 1/2/3/4^1,35–39^; ***iii*)** molecular chaperones, that recognize aberrantly exposed areas in out-of-shape and aggregating proteins, like αBcrystallin and Clusterin^40^. (Table S1,S11, Materials and Methods). RGS L106R and WT were screened for binding the array. We discovered eighteen peptides that bound selectively only to the mutant but not to the wild type protein (Fig. 2; Table 1).

**Table 1.**
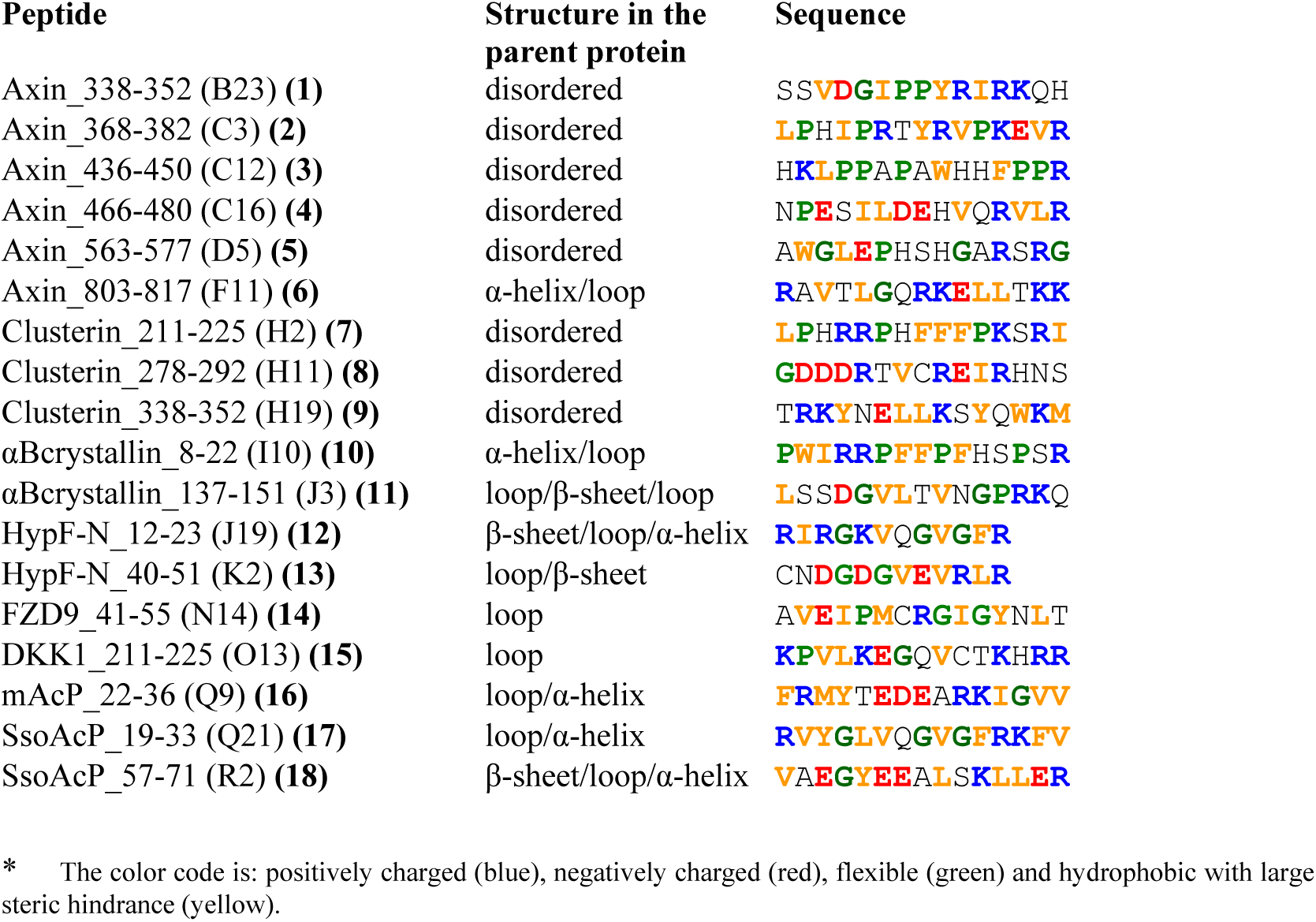
Peptides that bound specifically to RGS L106R in the peptide array*

**Figure 2.**
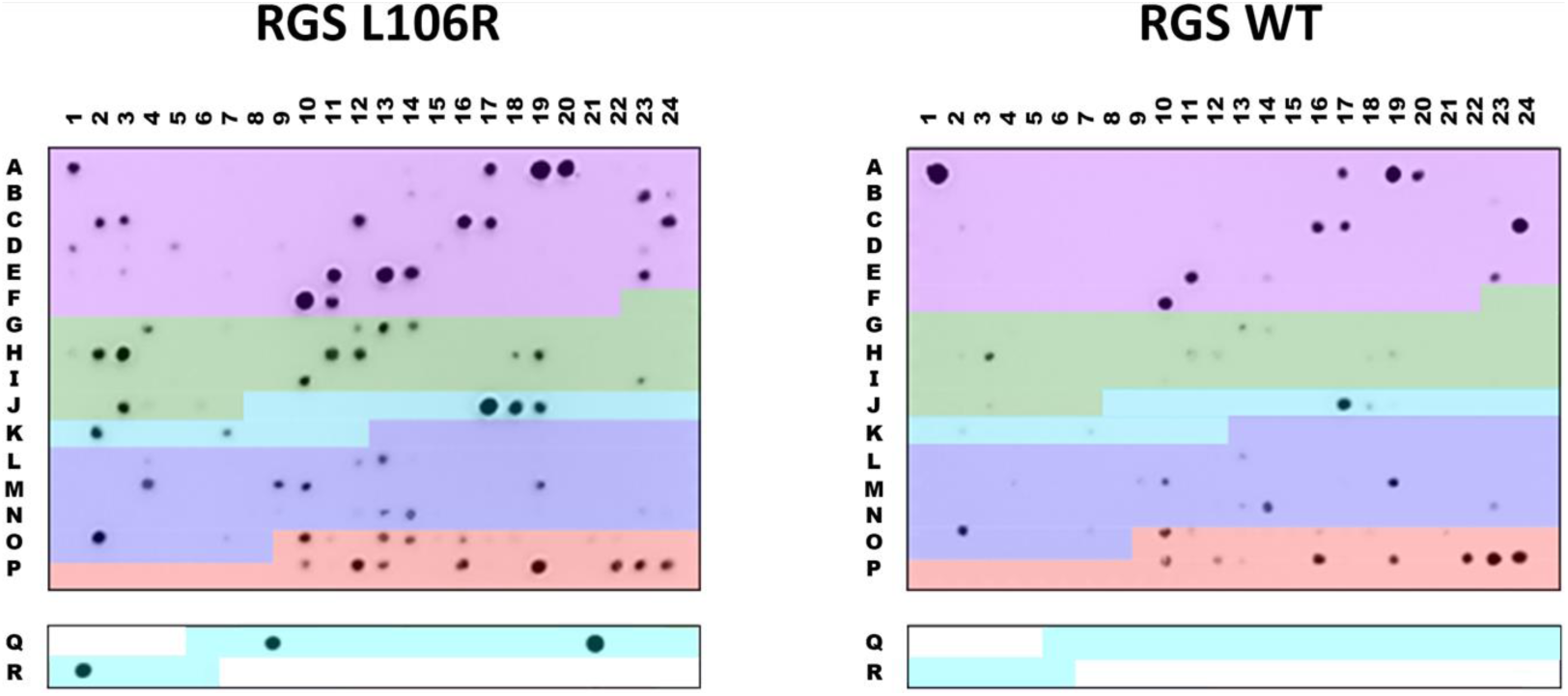
Peptide array screening identified specific binders of the aggregation-prone mutant L106R. Shown is the binding of RGS L106R (left) and RGS WT (right) to identical peptide arrays. The peptides are derived from the sequences of Axin (purple), αBcrystallin and Clusterin (green), Aβ_42_, HypF-N, SsoAcP and mAcP (light blue), Frizzled (blue) and DKK (pink). The array includes 15 or 12 residue peptides with an overlap of 8 or 7 residues. See supporting information for full peptide sequences.

### Selected peptides from the array inhibited amorphous aggregation

Next, we aimed to screen the RGSL106R binding peptides for their ability to prevent aggregation of RGSL106R in solution. To test this effect, we synthesized the eighteen peptides that bound selectively to RGS L106R on the peptide array and tested them for their ability to inhibit the established amorphous aggregation of this protein. The kinetics of RGS L106R aggregation was monitored by ANS, a fluorescent probe whose emission at 470 nm increases upon binding to hydrophobic patches^41^. The inhibitory effect of the peptides was quantified by normalizing the rate constant of the ANS time course in their presence (*k*_ANS,pep_) by that in their absence (*k*_ANS,ref_) (see Materials and Methods). The values of the normalized rate constant (*k’*_ANS_=*k*_ANS,pep_/*k*_ANS,ref_) represent the ability of the peptides to inhibit aggregation, since the precursor states are in this case stabilized by the peptides and large hydrophobic patches are solvent-exposed and free to bind ANS, but unable to propagate aggregation.

We then extracted the constant of *k’*_ANS_ dependence on peptide concentration to compare the inhibitory activity of the peptides (see Materials and Methods). A steeper dependence meant that a lower amount of peptide was needed for stabilizing the precursor and hence a more effective inhibition. Two out of the eighteen binding peptides, SsoAcP_19-33 **(17)** and αBcrystallin_8-22 **(10)**, almost completely inhibited RGS L106R aggregation in a dose-dependent manner (Fig. 3A,B; Table 2), with αBcrystallin_8-22 being a more effective inhibitor (Fig. 3C;Table 2). Consistently, TEM imaging of RGS L106R incubated in the presence of the two peptides revealed that αBcrystallin_8-22 suppressed aggregation at 400 μM, corresponding to 1:50 RGS:peptide molar ratio, while SsoAcP_19-33 only partly inhibited it at the same conditions (Fig. 3D-F; Table 2). All other RGS L106R-specific binders did not inhibit RGS L106R aggregation (Fig. S1; SsoAcP_57-71 is shown as representative for all).

**Table 2.**
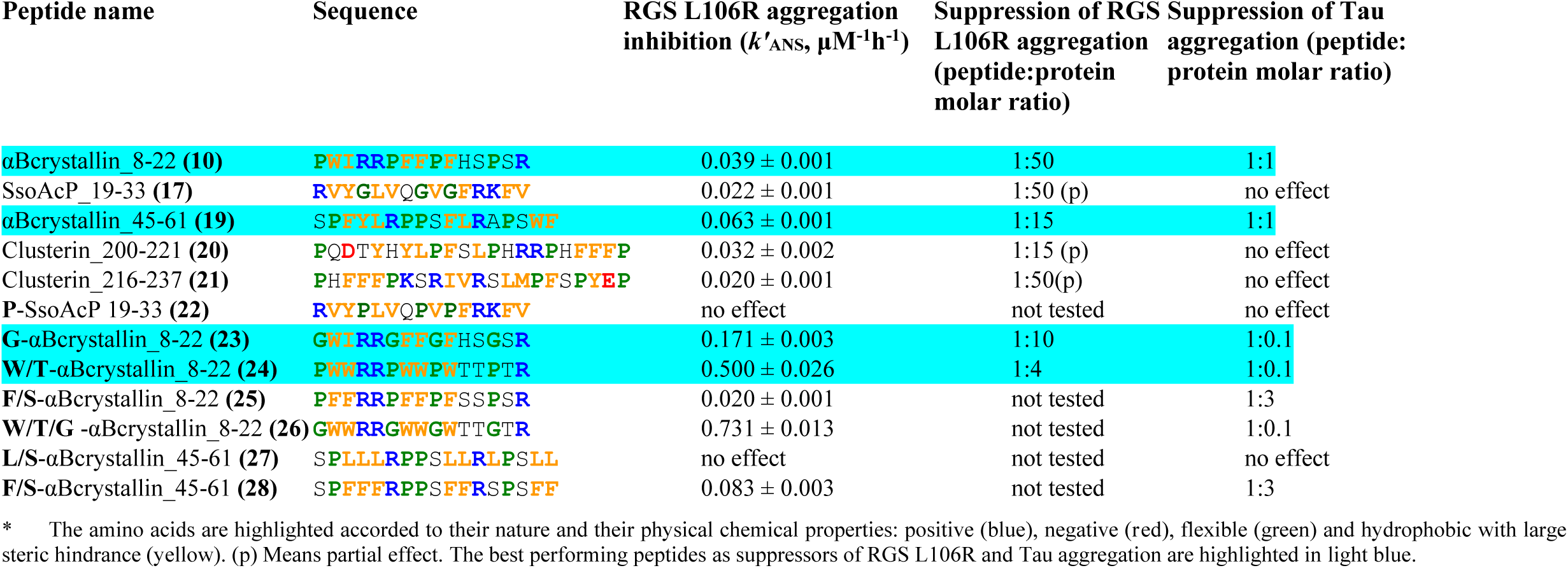
Activity of the designed inhibitors on protein aggregation*.

**Figure 3.**
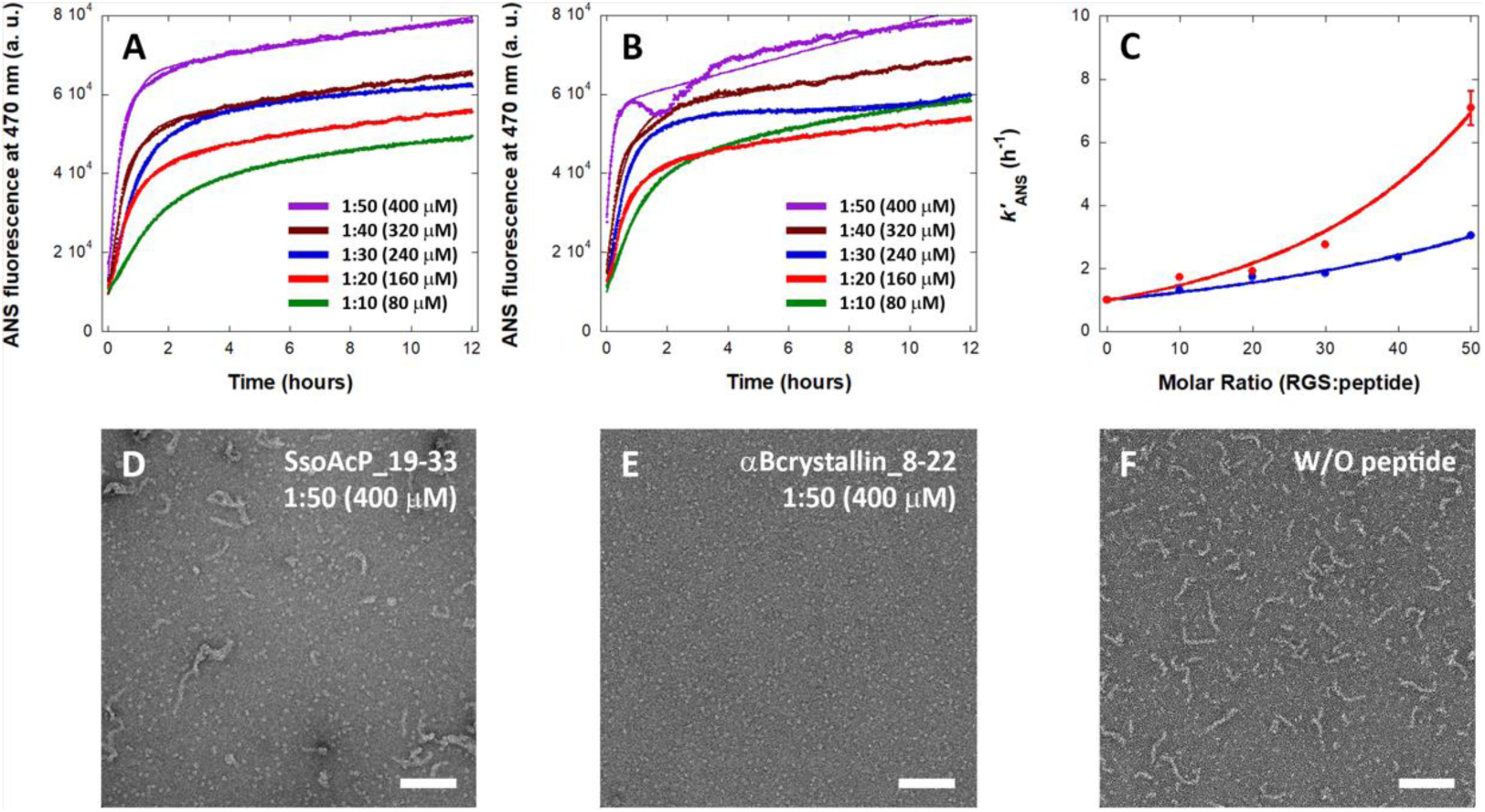
Two selected peptides inhibit RGS L106R aggregation. Shown are ANS time course kinetics of RGS L106R aggregation in the presence of different RGS:peptide molar ratios of A) SsoAcP_19-31 and B) αBcrystallin_8-22. C) Dose-dependence curve of *k’*_ANS_ on SsoAcP_19-31 (blue) and αBcrystallin_8-22 (red) concentration, indicating the inhibitory activity. Shown are also representative TEM image of RGS L106R incubated D, E) in the presence of the peptides or F) in their absence. The scale bar equals 100 nm in all cases.

### The molecular determinants are responsible for the anti-aggregation activity of the peptides

To understand the molecular principles that allow only some of the binders to inhibit aggregation, we searched for a pattern that is shared by the two inhibitors but is absent in the other binders. Each binder was analysed for its amino acid composition, hydrophobic content and residue distribution (see Materials and Methods). The sequences of SsoAcP_19-33 and αBcrystallin_8-22 were characterized by a distinct composition, being enriched in average by 3 aromatic and 2 flexible residues compared to a non-inhibitory binder (Fig. 4A,S2A; Tables S2,S4). Conversely, the content in aliphatic residues was similar in the two groups of peptides (Fig. 4A,S2A; Tables S2,S4). Although displaying similar amounts of positively charged residues as non-inhibitory binders, the inhibitory binders were enriched in arginine and depleted in lysine (Fig. 4A,S2A; Tables S2,S4,S10). Finally, the inhibitory binders were markedly less hydrophilic, reaching similar values for both hydrophobicity and hydrophilicity, at the brink of hydrophobic neutrality (Fig. 4B,S2B; Tables S3,S5). The residues of all the classes were evenly distributed throughout SsoAcP_19-33 and αBcrystallin_8-22 sequences without forming clusters. We tested whether such an even distribution was indispensable for activity and synthesized the Clusterin_211-225 peptide, which has a similar composition and hydrophobic content but presenting the aromatic residues clustered at mid-sequence. This peptide bound RGS L106R without inhibiting its aggregation (Table S2). To conclude, the molecular determinants required for a peptide to inhibit aggregation of RGS L106Rwere defined as: ***i*)** composition of 20-30% flexible, 30-40% aliphatic and 20-30% aromatic residues; ***ii*)** similar hydrophobicity and hydrophilicity; ***iii*)** even distribution of the amino acids according to their classes.

**Figure 4.**
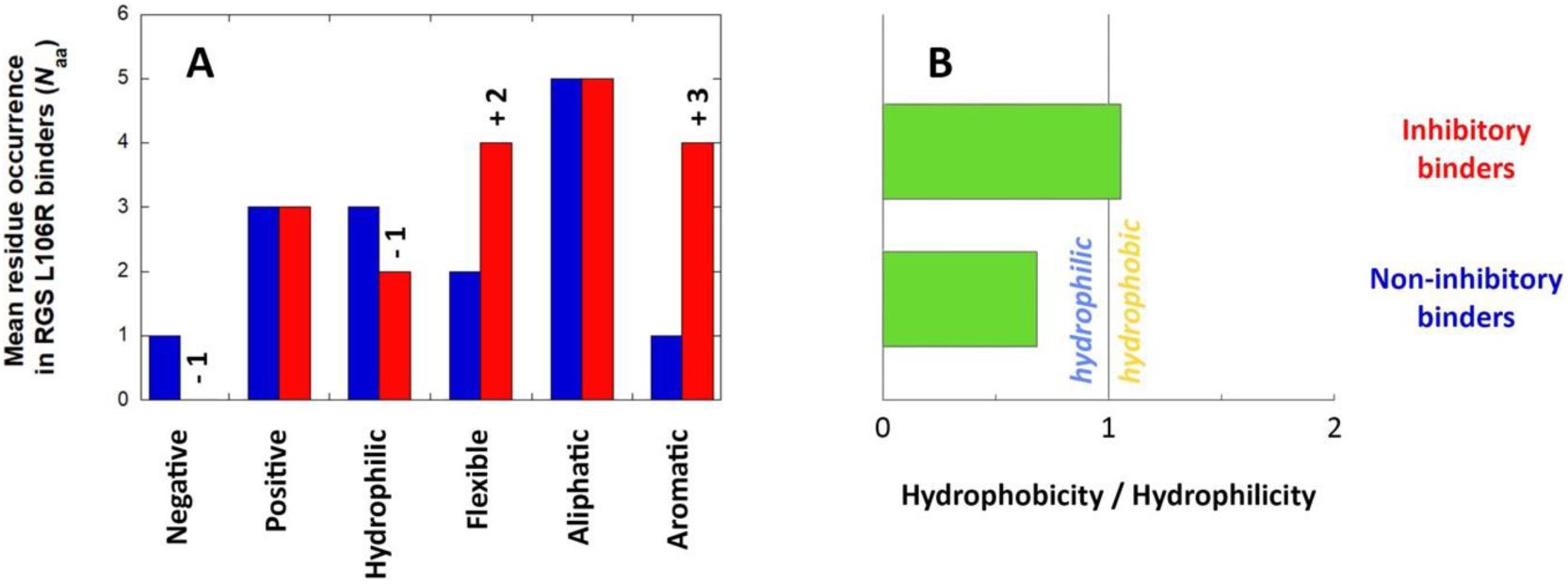
Analysis of the sequence of RGS L106R peptide binders in terms of composition and hydrophobicity. A) Mean residue occurrence by class of amino acids in inhibitory (red) and non-inhibitory (blue) binders expressed as number of amino acids (*N*_aa_). The relative enrichment or depletion is indicated on top of the column of the inhibitory binders. B) Hydrophobic content within inhibitory and non-inhibitory binders represented as ratio between total hydrophilicity (Hc_*N*_) and hydrophobicity (Hb_*N*_) calculated with the Roseman scale of hydrophobicity and normalised by the number (*N*) of residues of either pool of peptides (see Materials and Methods).

### The molecular determinants alone are responsible for the anti-aggregation activity

To further validate that the molecular determinants alone grant inhibitory activity to a peptide, we searched for unrelated protein sequences exhibiting the same signature motifs. We chose chaperone sequences, speculating that substrate binding clefts may use similar binding principles. Thus, we tested peptides derived from αBcrystallin and Clusterin. αBcrystallin_45-61 **(19)**, Clusterin_200-221 **(20)** and Clusterin_216-237 **(21)** displayed the required properties in terms of amino acid composition, hydrophobic content and distribution (Table S6,S7). Remarkably, these three peptides inhibited RGS L106R aggregation, with αBcrystallin_45-61 being the most potent inhibitor (Fig. 5A,B,6; Table 2). Clusterin_200-221 and Clusterin_216-237 also exerted an inhibitory effect but were less effective, as the decrease in aggregation was extensive but not complete (Fig. S3,6; Table 2).

**Figure 5.**
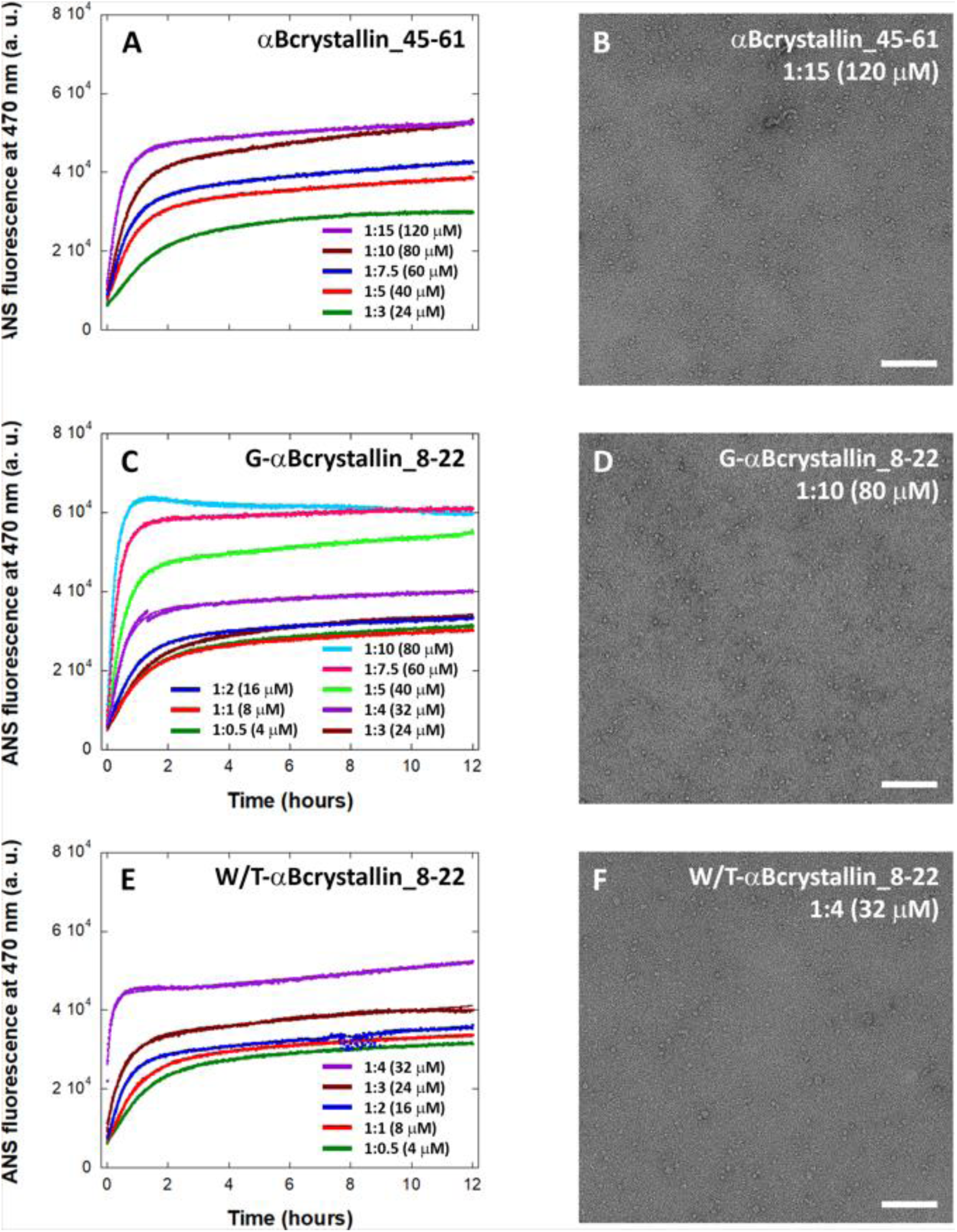
Designed peptides further suppress aggregation. A) ANS aggregation kinetics of RGS L106R in the presence of different concentrations of αBcrystallin_45-61 and B) representative TEM image of RGS L106R incubated with αBcrystallin_45-61. C) ANS aggregation kinetics of RGS L106R in the presence of different concentrations of **G-**αBcrystallin_8-22, and D) representative TEM images of RGS L106R incubated with **G-**αBcrystallin_8-22. E) ANS aggregation kinetics of RGS L106R in the presence of different concentrations of **W/T-**αBcrystallin_8-22, and F) representative TEM images of RGS L106R incubated with **W/T-**αBcrystallin_8-22. The scale bar in TEM images equals 100 nm.

**Figure 6.**
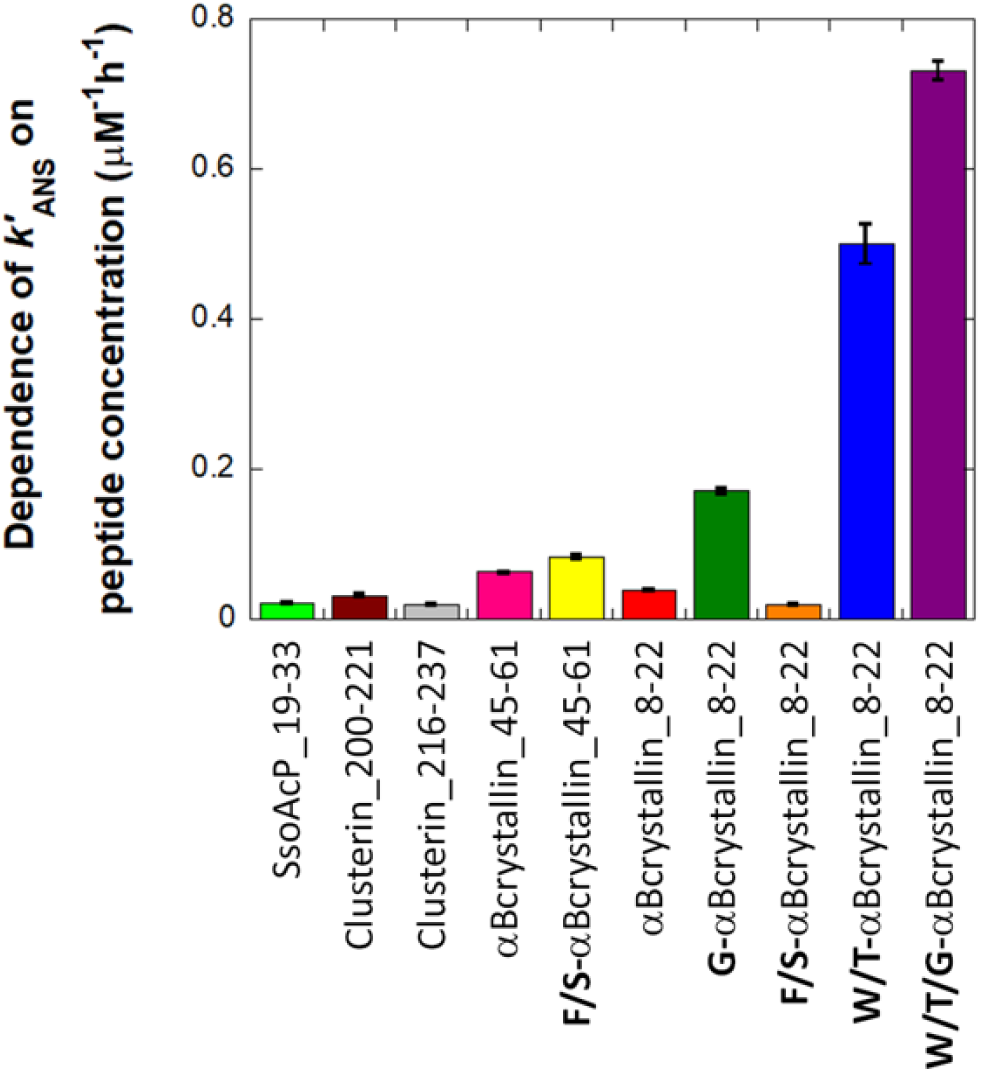
Dependence of *k’*_ANS_ on the concentration of the peptides. The result of the fit of the dose-dependence curve is plotted as indicative of the strength of the inhibitors.

### Optimizing flexibility of the peptide inhibitors

The next step was rationally designing a systematic mutational strategy for optimizing the molecular determinants and enhance the inhibitory activity (Table 2,S8,S9). First, we modulated the flexibility of the peptide chain to improve its adaptation to the variable interface of an aggregation precursor like the molten globular RGS L106R protein. Proline and glycine were mutually substituted since proline decreases the flexibility while glycine increases flexibility of the peptide chain. The mutations were performed on SsoAcP_19-33, the only peptide to bear glycine residues, and αBcrystallin_8-22, as a representative of all the other peptides that contain proline residues only. Gly-to-Pro substitutions (**P-**SsoAcP_19-33, **22**) suppressed the anti-aggregation activity of SsoAcP_19-33 (Fig. S4; Table 2). Conversely, Pro-to-Gly substitutions (**G-**αBcrystallin_8-22, **23**) increased the inhibitory activity of the peptide, which now completely suppressed RGS L106R aggregation at 80 μM, a 5-fold lower concentration than αBcrystallin_8-22 (Fig. 5C,D,6; Table 2). Such a stronger inhibitory activity in glycine-containing peptides suggested that glycine enhanced inhibition better than proline, irrespective of the sequence of the peptide. A higher degree of internal freedom was hence beneficial for a peptide molecule to recognize and bind the fluctuant structural conformations of an aggregation precursor state.

### Optimizing the aromacity of the peptide inhibitors by tryptophan residues

We then fine-tuned the hydrophobicity of the two peptides that displayed the strongest inhibitory activity: αBcrystallin_8-22 and αBcrystallin_45-61. Modifying the amount and the density of hydrophobicity throughout the peptide sequence is expected to modulate the ability to recognize the exposed hydrophobic patches that trigger aggregation. First, variants of the peptides were designed to maintain the same total hydrophobicity as the parent peptides. The hydrophobic density was equalized throughout the sequence by substituting all residues of the same type by one specific residue with the same property (Table S8,S9). In αBcrystallin_8-22, we substituted the hydrophobic residues by tryptophan and the hydrophilic ones by threonine, while in αBcrystallin_45-61 we substituted by leucine and serine (**W/T-** αBcrystallin_8-22, **24**; **L/S-**αBcrystallin_45-61, **27**) (Table 2). **W/T-**αBcrystallin_8-22 was 13-fold more active than αBcrystallin_8-22 and 3-fold more active than of **G-**αBcrystallin_8-22 (Fig. 5E,6; Table 2). Consistently, the complete inhibition of RGS L106R aggregation was observed with TEM imaging at **W/T-**αBcrystallin_8-22 concentrations as low as 32 μM or 1:4 RGS:peptide molar ratio, a 12.5-fold lower concentration than αBcrystallin_8-22 (Fig. 5F). On the contrary, **L/S-**αBcrystallin_45-61 did not show any inhibitory activity (Fig. S5A, Table 2).

We designed further variants of αBcrystallin_8-22 and αBcrystallin_45-61 with maximized hydrophobic density, while maintaining fixed the total hydrophobicity (Table S8,S9). The hydrophobic residues were substituted with phenylalanine and the hydrophilic residues with serine to compensate for the increase in hydrophobicity (**F/S-**αBcrystallin_8-22, **25**; **F/S-**αBcrystallin_45-61, **28**) (Table 2). In the case of **F/S-**αBcrystallin_45-61, the substitutions resulted in a 1.5-fold increase in the inhibitory activity compared to the original peptide, while for **F/S-**αBcrystallin_8-22 it decreased to about half that of the non-substituted peptide (Fig. S5B,C; Table 2). This means that tryptophan contributes to the inhibition much more than phenylalanine.

### Optimization of the inhibitors reveals a new molecular basis for inhibition

The analysis of the relative amino acid abundance revealed that phenylalanine, valine, proline and arginine are strongly enriched in the natural sequence of the inhibitors as compared to binders (Table S10). Conversely, tryptophan and glycine, that perform as best enhancers of inhibitory activity, are only slightly enriched (Table S10). Therefore, the optimization identified beneficial sequence elements that were not naturally encoded in the original peptide set. Based on this, tryptophan, threonine and glycine substitutions were incorporated together in the sequence. The resulting peptide (**W/T/G-** αBcrystallin_8-22, **26**) had an inhibitory activity that is slightly higher than the sum of the values observed for **G-**αBcrystallin_8-22 and **W/T-**αBcrystallin_8-22. This suggests that the effects of optimized hydrophobicity and flexibility are additive (Table 2; Fig. S5D,6).

### The designed peptides inhibit Tau amyloid aggregation

We then assessed whether our method of discovery and optimization of inhibitors is general and applicable to proteins aggregating via different mechanisms. The chaperone-derived peptides were tested as inhibitors of Tau aggregation. Tau aggregates into fibrils of defined, regular structure in Alzheimer’s and other Tauopathies^42,43^. The αBcrystallin-derived peptides that inhibited RGS L106R aggregation also inhibited the aggregation of 20 μM Tau microtubule binding region (Q244-E372), an aggressively aggregating Tau fragment causing neurodegeneration in mice, but with higher efficiency^44^. αBcrystallin_8-22 and αBcrystallin_45-61 suppressed Tau aggregation at the equimolar concentration of 20 Mm (Fig. 7A; Table 2). The optimized peptides **G-**αBcrystallin_8-22, **W/T-**αBcrystallin_8-22 and **W/T/G-** αBcrystallin_8-22 potently inhibited Tau aggregation at the substoichiometric concentration of 2 μM (Fig. 7B; Table 2). **F/S-**αBcrystallin_8-22 and **F/S-**αBcrystallin_45-61 also inhibited Tau aggregation but at superstoichiometric concentrations, while **L/S-**αBcrystallin_45-61 did not exert any inhibition (Table 2). This suggests that the molecular determinants grant anti-aggregation activity towards different kinds of protein aggregation according to the same principles.

**Figure 7.**
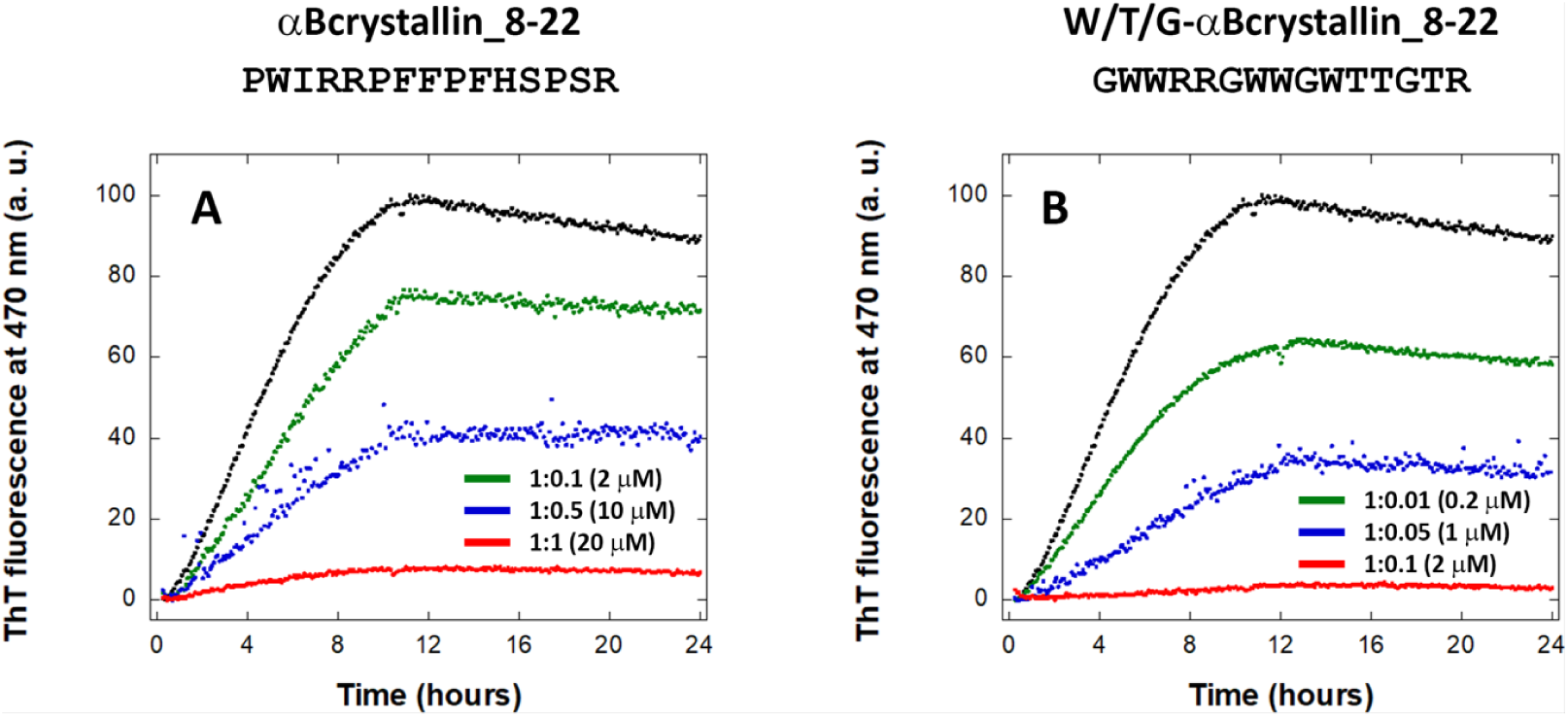
Aggregation of Tau microtubule binding region (Q244-E372) in the presence of different concentrations of A) αBcrystallin_8-22 and B) **W/T/G-**αBcrystallin_8-22. The black trace is the protein in the absence of peptide.

## Discussion

### Molecular determinants for designing aggregation inhibitors

In this work we present a new method for designing peptide inhibitors against protein aggregation based on the idea that the inhibitory activity can be an intrinsic property of an amino acid sequence. We obtained the molecular determinants of anti-aggregation activity by combining the output of two assays: one for binding (peptide array) and the other for activity (ANS fluorescence) against the aggregation of Axin RGS L106R, here used as a model protein for early aggregative interactions. As a molten globule, RGS L106R allowed to screen selectively for inhibitors of the aggregation precursor state, without being specific for a mechanism.

Comparing the inhibitory and non-inhibitory binding peptide sequences revealed unique molecular determinants, that were used to design new inhibitors. The inhibitors were characterized by increased content of aromatic and flexible residues, evenly spread over the sequence. A design strategy based on the rational optimization of the molecular determinants produced peptides with high inhibitory activity and broad recognition abilities towards amorphous and fibrillar aggregation, reminiscent of that of molecular chaperones. Among these, **W/T-**αBcrystallin_8-22 (PWWRRPWWPWTTPTR) was obtained as an optimized version of the highly versatile αBcrystallin_8-22 peptide, and stood out as the best lead compound for further development in drug discovery.

### The molecular interactions responsible for the cross-mechanism inhibitory activity

Proline is an amyloid-inhibiting residue thanks to its β-breaker abilities^45^. However, our mutational campaign revealed that glycine enhances inhibition more effectively. With a wider distribution of dihedral angles, glycine may adjust better the peptide backbone to the structurally fluctuating binding sites of the molten globular RGS L106R and the disordered Tau^46^. Conversely, tryptophan enhances inhibition thanks to the π-stacking and H-bonding capabilities of its side chain. Hydrophobicity does not confer the inhibitory activity *per se*, as leucine substitutions suppressed the inhibition and phenylalanine substitutions left the inhibitory potency unchanged. This means that the π-stacking interactions, which exist in both Phe and Trp but not in Leu, are key for the peptide activity, but not enough for its improvement. The combined π-stacking and H-bonding interactions may grant tryptophan the ability to recognize mixed hydrophobic/hydrophilic surfaces and anchor firmly to aggregating interfaces, allowing for multi-targeting. Consistently, a solvent exposed tryptophan is directly involved in the recognition of different amyloid strands in WW2 peptides-mediated inhibition^16^. Naphthoquinone-tryptophan derivatives inhibit a number of amyloidogenic processes and small molecules bearing indole building blocks are potent and general inhibitors for amyloids^47,48^. Similar considerations are made for arginine, the only charged residue found in all peptides that shares with tryptophan the mixed nature of π-system and hub of H-bonding. RGSL106R hotspot is enriched in aromatic residues (F117, W118, F119, F124) and is electrostatically neutral (D113, D116, R125, K126), so arginine likely contributes to binding via π-stacking and H-bonding. This is consistent with the finding that the disordered interactome of fibrillar Tau is enriched in arginine, meaning that such residue is key to recognize and inhibit the aggregating protein^49^.

Together, our data suggest that a high density of both π-stacking and H-bonding is required to inhibit protein aggregation as a whole, irrespective of sequence, structure or mechanism (Fig. 8A). The absence of clusters of residues with similar properties throughout the sequence suggests that concentrating flexibility, π-stacking or H-bonding in distinct spots of the molecule is not advantageous for inhibition. Hence, an evenly distributed high density, instead of peaks of density, grants and boosts inhibitory activity towards aggregating proteins (Fig. 8B).

**Figure 8.**
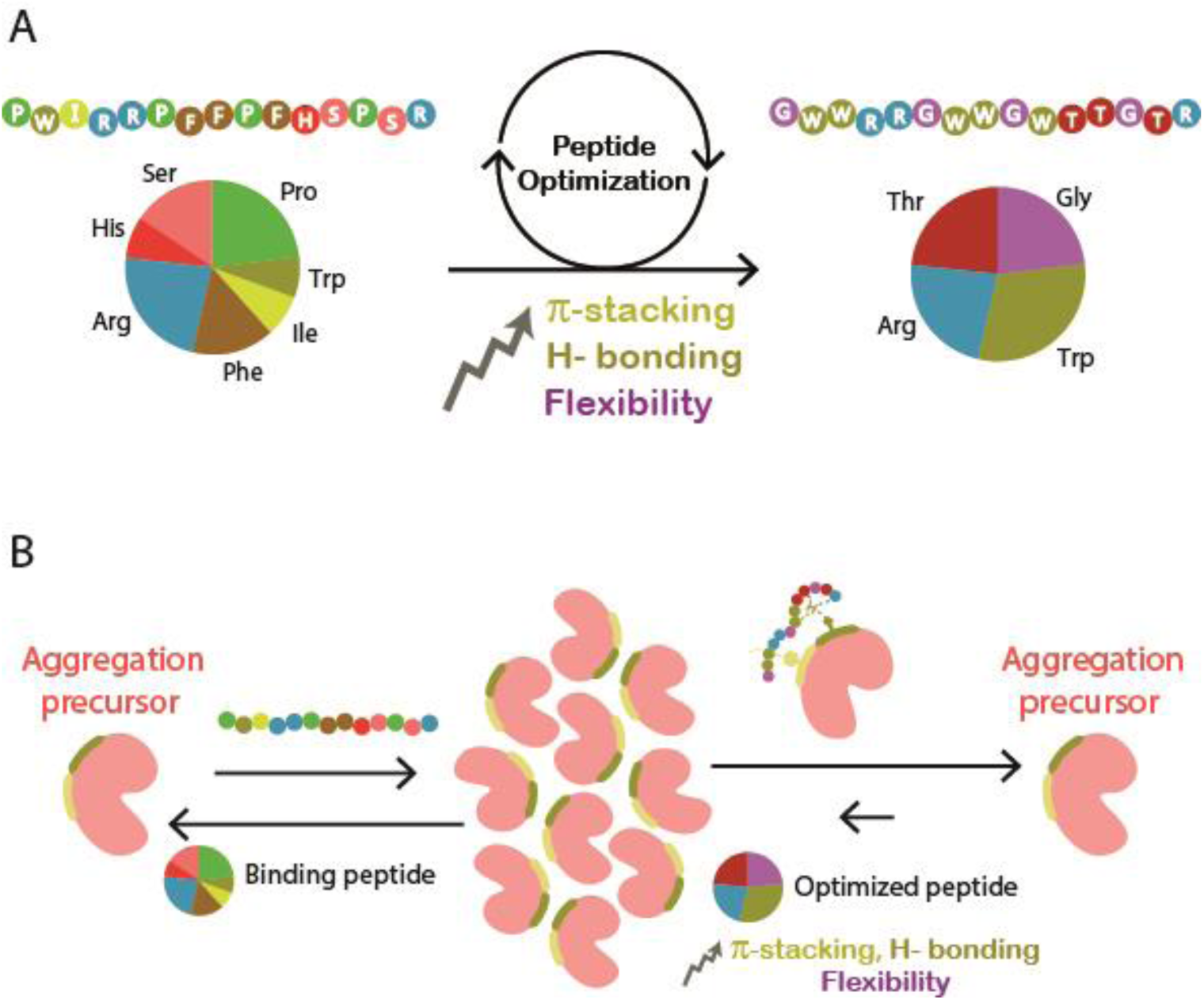
Properties and activity of an optimized peptide aggregation inhibitor. A) The sequence of αBcrystallin_8-22 is shown as a scheme on the left and the optimized peptide sequence (W/T/G αBcrystallin_8-22) on the right. The optimized sequence includes increased distribution of flexible (purple) and hydrophobic (yellow) residues. The distribution of positively charged (blue), and hydrophilic (pink) residues did not change. A composition of 27% glycine, 33% tryptophan, 20% arginine residues evenly distributed throughout the sequence grants the backbone flexibility and the high density of H-bonding and π-stacking interactions needed for inhibitory activity. B) The optimized peptide (right panel) shifts the equilibrium between aggregation to the precursor state more dramatically than the unoptimized binding peptide (right panel).

### Advantages over other peptide targeting strategies

Our peptides proved inherently multi-targeting and suppressed Tau and RGS L106R aggregation alike. As such, they represent an alternative to the current sequence-based strategies that ensure a highly specific recognition by designing complementary β-sheets^7,11^. Currently there are no effective inhibitors in the clinic, and we present a new strategy that can be further exploited for drug design. Similarly, p53 aggregation is not inhibited by targeting one hotspot at a time, and the sequence-based ReACp53 peptide releases native p53 incorporated in the growing aggregate upon fibril disassembly, without stabilizing all hotspots of the aggregating protein^14,50,51^. The sequence-based approach for inhibition hence does not ensure a full, reliable and long-lasting protection against protein aggregation. Using the molecular determinants to design inhibitors overcomes all the limitations of taking an aggregation hotspot as a “cast”. Being intrinsic inhibitors, the determinants-bearing peptides are not specific for any mechanism, protein or strand, nor need to match the structure/sequence requirements of any aggregation hotspot to gain activity. Contrary to inhibitors designed by other approaches, they can neutralize several aggregation hotspots of one or more proteins yielding a complete inhibition of aggregation, and can be improved through mutational screening.

### Implications for drug design

Based on their ability of inhibiting cancer and neurodegeneration -related aggregates, our peptides are ideal cross-therapeutics for both pathologies^52^. They may inhibit nano-aggregation of Axin RGS mutants *in vivo*, restoring the Wnt suppressive function and inducing the regression of the related cancer phenotypes^31^. They may also act as lead compounds in the drug discovery against the neurodegenerative diseases having Tau aggregation as molecular etiology^33^. Being inherently multi-targeting, chaperone-like peptides may intercept the aggregation precursor state of other proteins, enlarging the spectrum of their direct therapeutic applicability. All together they constitute an ensemble of molecules with basal anti-aggregation activity for high-throughput approaches and screenings, amenable to grafting onto engineered chaperones or antibodies to potentiate their versatility and efficacy as protein therapies. The design of chaperone-like peptides hence constitutes a promising direction *per se* and a great advance in the existing methods for drug discovery.

## Acknowledgements

We thank Dr. Yael Levi-Kalisman of the Center for Nanoscience and Nanotechnology at the Hebrew University of Jerusalem. This work was supported by the Innovative Training Network 608180 “WntsApp” within by Marie-Curie Actions of the 7th Framework program of the EU. AF thanks The Minerva Center for Bio-Hybrid complex systems and the Saerree K. and Louis P. Fiedler Chair in Chemistry.S.G.D.R was supported by Campaign Team Huntington, Alzheimer Nederland and a ZonMW TOP grant (“Chaperoning axonal transport in neurodegenerative disease”; No. 91215084).

## Materials and methods

### Expression and purification of RGS WT, RGS L106R and Tau

For the peptide array assays, RGS WT and RGS L106R protein constructs comprising two His and one NUS tags (His-NUS-His-RGS WT and His-NUS-His-RGS L106R) were expressed as reported^31^ and purified as follows. The bacteria were lysed with a microfluidizer (Microfluidics) and the lysate separated from the pellet by centrifuging at 15000 rpm for 45 minutes. The soluble fraction was purified by affinity chromatography using an ÄKTA Explorer FPLC system (GE Healthcare) loaded with a Nickel Sepharose FF 8 mL resin column. Elution was performed with an imidzole gradient in 50 mM tris HCl pH 7.4, 150 mM NaCl, 5 mM β-mercaptoethanol. The fused His-NUS-His tag was obtained by cleavage with 2.2 μM TEV protease upon overnight incubation at 4 °C under mild agitation. The protein constructs were further purified by size exclusion chromatography using an ÄKTA Explorer FPLC system (GE Healthcare) loaded with two Superdex75 200 mL columns. The purity of the protein was confirmed by electrophoresis with SDS-page gels. The protein constructs were concentrated with Vivaspin 20 5000molecular weight cut-off centrifuge tubes (GE Healthcare) and stored at -80 °C. For all other experiments, the RGS L106R protein was expressed and purified as reported^32^. Tau was expressed and purified as described^53^.

### Peptide array screening

The CelluSpot™ peptide micro-arrays were synthesized by INTAVIS Bioanalytical Instruments AG, Köln, Germany. The peptides were acetylated at their N-termini and attached to a cellulose membrane via their C-termini through an amide bond. The arrays were incubated for 3 hours with 5% (w/v) skimmed milk to 50 mM Tris HCl pH 7.5, 150 mM NaCl, 0.05% (v/v) Tween20 as blocking solution. The arrays were washed three times with buffer and incubated overnight in the presence of 4 μM His-NUS-His, His-NUS-His-RGS WT or His-NUS-His-RGS L106R constructs at 4 °C in blocking solution. The arrays were washed three times with buffer and incubated with 5 μM of a monoclonal antibody anti-His HRP-conjugated for 1 hour at room temperature. The arrays were washed three times with buffer and immunodetected by chemoluminescence using ECL reagents and a Vilber Fusion FX camera.

### Peptide synthesis and purification

The peptides were synthesized with a Liberty Microwave Assisted Peptide Synthesizer (CEM, Matthews, NC, USA) using standard 9-fluorenylmethoxycarbonyl (Fmoc) chemistry, N,N’-diisopropylcarbodimide (DIC)/oxima as coupling reagents and a Rink-Amide resin with 0.546 mmol/g substitution. The peptides were cleaved from the resin with a mixture of 95% (v/v) trifluoroacetic acid (TFA), 2.5% (v/v) triisopropylsilane (TIS), 2.5% (v/v) triple distilled water (TDW) agitating vigorously for 3 hours at room temperature. The volume was decreased by N_2_flux and the peptides precipitated by addition of 4 volumes of diethylether at -20 °C. The peptides were sedimented at -20 °C for 30 minutes, then centrifuged and the diethylether discarded. The peptides were washed three times with diethylether and dried by gentle N_2_ flux. The solid was dissolved in 1:2 volume ratio of acetonitrile (ACN):TDW, frozen in liquid Nitrogen and lyophilized. The peptides were purified on a Jasco LG2080 HPLC using a reverse-phase C18 preparative column with a gradient comprised between 10 and 40% (v/v) ACN:TDW at a rate of 0.5%/min. The peptides were identified by ESI mass spectrometry (MS) performed on an LCQ Fleet Ion Trap (Thermo Scientific) mass spectrometer. Peptide masses were calculated from the experimental mass to charge (m/z) ratios from the observed multiple-charged species of a peptide. Deconvolution of the experimental MS data was performed using the MagTran v1.03 software. The purity of the peptides was checked using a Merck Hitachi D7000 analytical HPLC and a reverse-phase C8 analytical column.

### UV spectroscopy

UV spectra were recorded with a Shimadzu UV-1650PC spectrophotometer using a quartz cuvette of 0.1 cm path length for far-UV spectroscopy. The extinction coefficient at 280 nm (ε_280_) of RGS WT and RGS L106R in TDW was 0.936 M^−1^cm^−1^, as computed by ExPASy. The extinction coefficient at 360 nm (ε_360_) of 8-Anilinonaphthalene-1-sulfonic acid (ANS)^54^ in TDW was 5700 M^−1^cm^−1^. The extinction coefficient at 412 nm (ε_412_) of Thioflavin T (ThT)^55^ in TDW was ε_412_ = 36000 M^−1^cm^−1^.

### ANS fluorescence

25 μL of 32 μM RGS L106R were added to 75 μL of aggregating solution in the absence or in the presence of the peptides at different concentrations. The solution was pipetted 10 times to obtain homogeneity. In all conditions, the solutions were 8 μM RGS L106R in 50 mM tris HCl pH 7.4, 150 mM NaCl, 5 mM β-mercaptoethanol, 24 μM ANS or ThT, 25°C. The aggregation kinetics were acquired with a BioTek Synergy H1 Hybrid Reader (Thermo Scientific) plate reader using 96 wells half-area plates (Costar 3696) for 100 μL volume solutions. ANS was excited at 380 nm and the emission collected at 470 nm. The time course of ANS traces were fit using a single exponential plus line equation (Eq. 1).

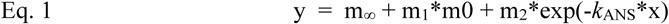

where m_∞_ is the ANS emission at the end of the kinetics, m_1_ is the angular coefficient of the linear correlation, m_2_ is the difference between m_∞_ and the ANS emission at the beginning of the kinetics, and *k*_ANS_ the rate constant of the exponential phase that was used to characterize the RGS L106R aggregation process^32^.

The rate constant in the presence of different concentrations of the peptides (*k*_ANS,pep_) was divided by that of the reference reaction in their absence (*k*_ANS,ref_) to yield a normalized rate constant (*k’*_ANS_=*k*_ANS,pep_/*k*_ANS,ref_). For each peptide, *k’*_ANS_ values at different concentrations were fit with an exponential rise equation (Eq. 2).

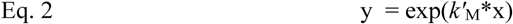

where *k’*_M_ is the correlation constant used to characterize the dose-dependence of the peptides inhibitory effect on RGS L106R aggregation. The non-linear fit was probably due to non-specific interplays of ANS with surface hydrophobicity.

### ThT fluorescence

Aggregation of 20 µM Tau-RD* (N-terminally FLAG-tagged (DYKDDDDK) Human Tau-RD (Q244-E372, also referred to as K18, with pro-aggregation mutation DK280) in aggregation buffer (25 mM HEPES-KOH pH 7.5, Complete Protease Inhibitors (1/2 tablet/50 ml), 75 mM KCl, 75 mM NaCl and 10 mM DTT) was stimulated by 5 µM heparin Low Molecular Weight in the presence of 60 µM ThioflavinT (Sigma-Aldrich). Aggregation was performed in transparent, lidded Greiner 96-well plates (Sigma-Aldrich). To assess the impact of peptides, samples were supplied as indicated in each experiment. Fluorescent spectra were recorded every 10 minutes for 24hours with a SpectraMax i3 (Molecular Devices). For each well, total volume was 100 µL.

### Transmission electron microscopy (TEM)

RGS L106R was incubated at a concentration of 8 μM in 50 mM tris HCl pH 7.4, 150 mM NaCl, 5 mM β-mercaptoethanol at 25°C for 12 hours. The aggregation reaction solution was diluted 10x in the same buffer prior to sample preparation. A drop of 3-5 μL sample was applied to a glow discharged TEM grid (carbon supported film on 300 mesh Cu grids, Ted Pella, Ltd.). After 30 sec the excess liquid was blotted, the grids were stained with 2% uranyl acetate for 30-60 sec, blotted and allowed to dry in air. The samples were examined using FEI Tecnai 12 G^2^ TWIN TEM operated at 120 kV. The images were recorded by a 4K × 4K FEI Eagle CCD camera.

### Sequence analysis

Amino acids were grouped into six classes according to their biophysical properties: positively charged, negatively charged, hydrophilic, flexible, aliphatic and aromatic. The amino acid composition was expressed as relative abundance and occurrence. The relative abundance was calculated as percent amount of residues of the aforementioned classes (Tables S2,S4,S6,S8). The occurrence was calculated as approximate number of amino acids of the aforementioned classes corresponding to relative abundance in a peptide 15 residues long (Tables S4). Both abundance and occurrence were determined for each individual peptide and as a mean of all inhibitory or non-inhibitory binders. The hydrophobic content was determined by summing up the Roseman hydrophobicity score (HS) of each residue to yield total hydrophobicity (Hb, positive values) and hydrophilicity indexes (Hc, negative values). The mean hydrophobicity (Hb_*N*_) and hydrophilicity per residue (Hc_*N*_) were obtained by normalising Hb and Hc for the number of residues present in each peptide (12 or 15 aa) and in all theinhibitory (30 aa) or non-inhibitory binders (264 aa) (Tables S3,S5,S7,S9).The distribution was defined as even or clustered, depending on whether the residues of the same class are scattered throughout the sequence or all close to one another.

