## Supporting information for "A peptide strategy for inhibiting different protein aggregation pathways in disease"

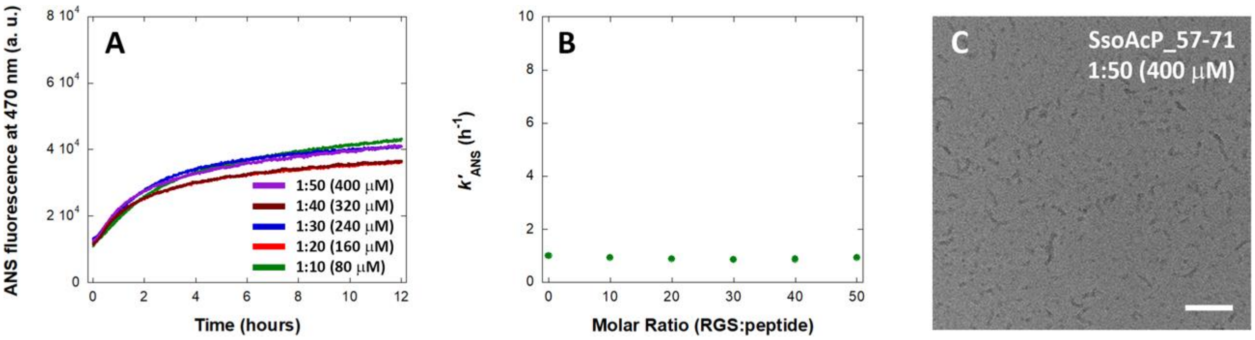


**Figure S1.** Effect of the non-inhibitory binders on RGS L106R aggregation. ANS time course aggregation kinetics of RGS L106R in the presence of different concentrations of SsoAcP_57-71. B) Dose-dependence of the $k'_{\mathrm{ANS}}$ of RGS L106R aggregation on SsoAcP_57-71. C) Representative TEM image of RGS L106R incubated in the presence of SsoAcP_57-71 (positive staining). The scale bar equals 100 nm.

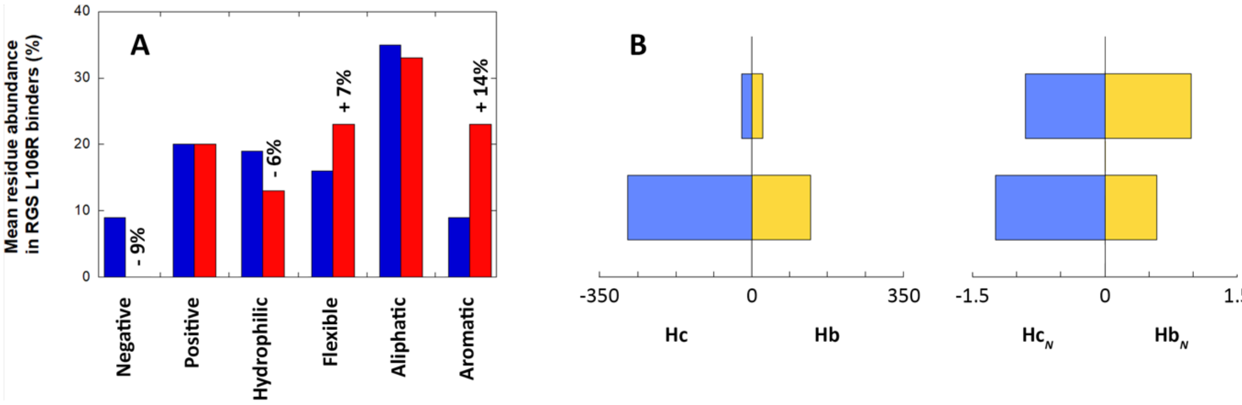


**Figure S2.** Analysis of the sequence of RGS L106R peptide binders in terms of composition and hydrophobicity. A) Mean residue abundance by class of amino acids in inhibitory (red) and non-inhibitory (blue) binders expressed as percent amount. The relative enrichment or depletion is indicated on top of the column of the inhibitory binders. B) Hydrophobic content within inhibitory and non-inhibitory binders represented as sum of hydrophilicity (Hc, light blue)

and hydrophobicity (Hb, yellow) scores calculated with the Roseman scale of hydrophobicity and normalized by number of residues (*N*) of either pool of peptides (Hc$_N$, light blue; Hb$_N$, yellow) (see Materials and Methods).

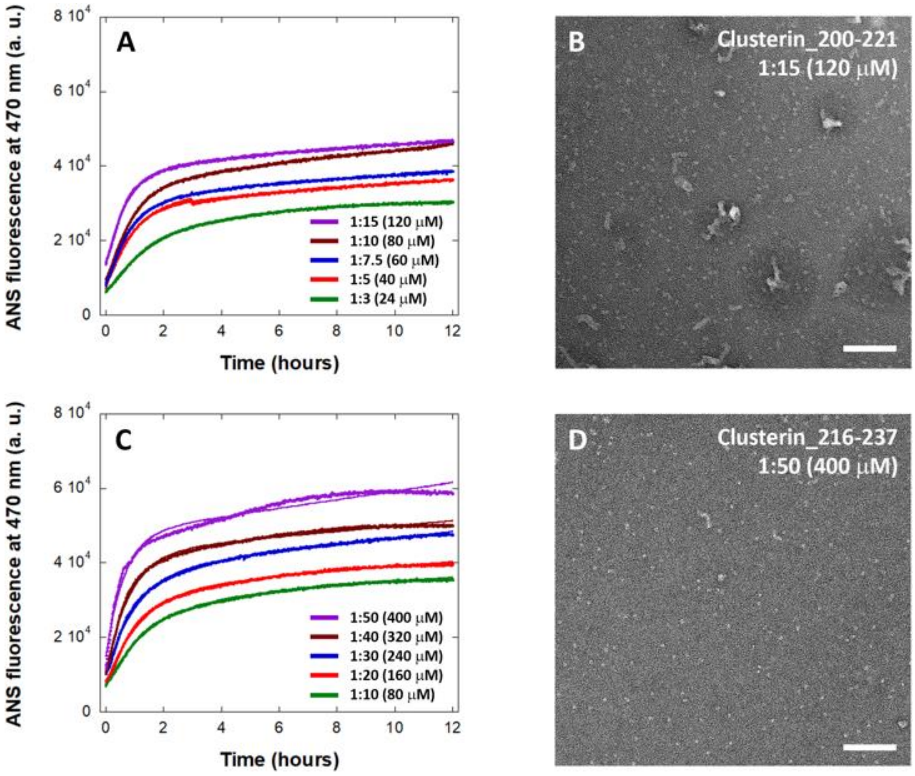


**Figure S3.** Effect of designed peptides on RGS L106R aggregation. A) ANS time course aggregation kinetics of RGS L106R in the presence of different concentrations of Clusterin_200-221. B) Representative TEM images of RGS L106R incubated in the presence of Clusterin_200-221. C) ANS time course aggregation kinetics of RGS L106R in the presence of different concentrations of Clusterin_216-237. D) Representative TEM images of RGS L106R incubated in the presence of different concentrations of Clusterin_216-237. The scale bar in TEM images equals 100 nm.

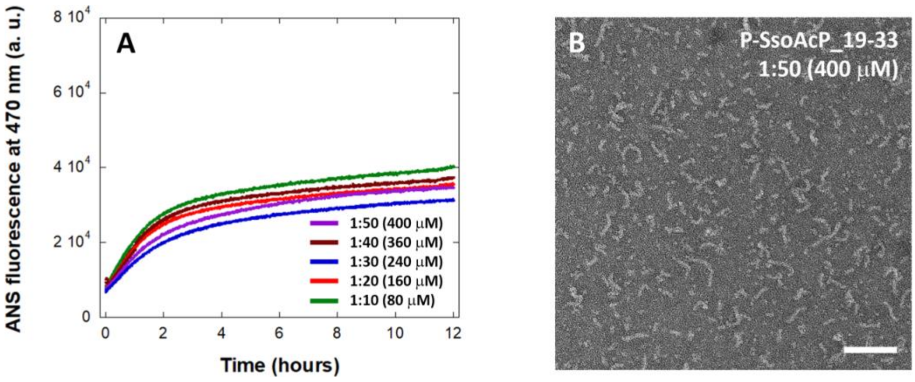


**Figure S4.** Effect of designed peptides on RGS L106R aggregation. A) ANS time course aggregation kinetics of RGS L106R in the presence of different concentrations of **P**-SsoAcP_19-33. B) Representative TEM images of RGS L106R incubated in the presence of **P**-SsoAcP_19-33. The scale bar equals 100 nm.

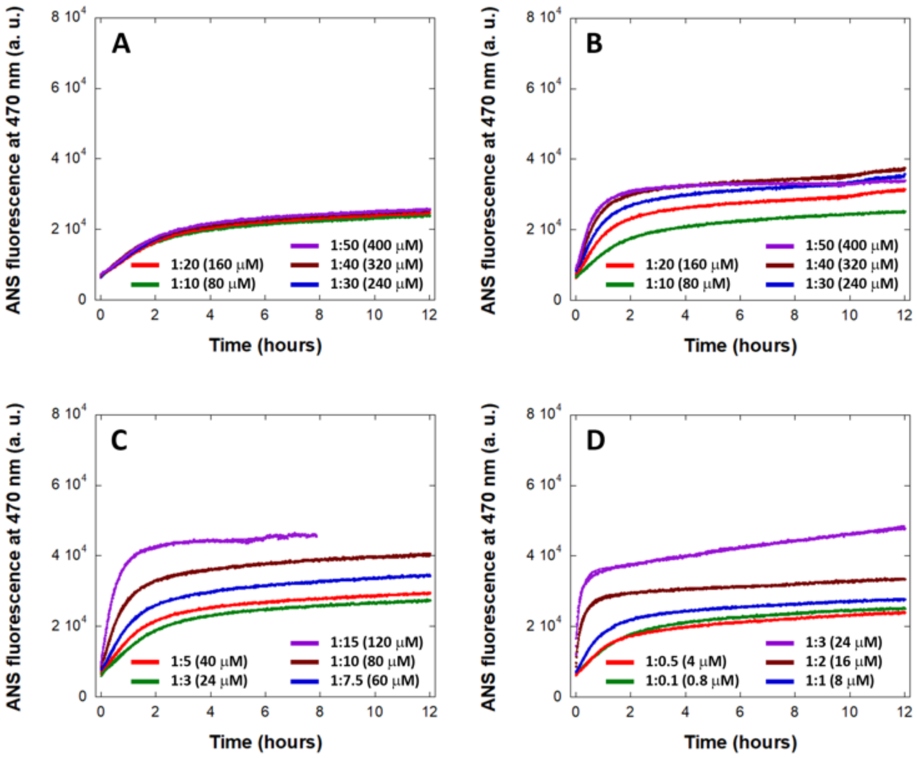


**Figure S5.** Effect of designed peptides on RGS L106R aggregation. ANS time course aggregation kinetics of RGS L106R in the presence of different concentrations of A) **L**-αBcrystallin_45-61, B) **FS-**αBcrystallin_8-22, C) **FS-**αBcrystallin_45-61, and **WTG-**αBcrystallin_8-22.

**Table S1. Proteins from which peptides included in the array were derived**

| Protein | Biological function |
|---|---|
| Axin 1 | Scaffold protein of the Wnt pathway. |
| αBcrystallin | Intracellular chaperone associating with amyloids. |
| Clusterin | Extracellular chaperone associating with amyloids. |
| $A\beta_{42}$ | Main component of amyloid plaques in Alzheimer's Disease. |
| mAcP | Model protein for amyloid aggregation via misfolded state. |
| SsoAcP | Model protein for amyloid aggregation via native-like state. |
| HypF-N | Model protein for amyloidmisfoldedoligomers. |
| CRD of Fzds | Receptors of the Wnt pathway. |
| CRD of DKKs | Extracellular antagonists of the Wnt pathway. |

**Table S2.** Relative abundance of amino acids in RGS L106R binders*.

| Peptide | Negative | Positive | Hydro-philic | Flexible | Aliphatic | Aromatic |
|---|---|---|---|---|---|---|
| Axin_338-352 **(1)** | 6.7 % | 20.0 % | 26.7 % | 20 % | 33.3% | 6.7 % |
| Axin_368-382 **(2)** | 6.7 % | 26.7% | 13.3 % | 20 % | 46.7 % | 6.7 % |
| Axin_436-450 **(3)** | - | 13.3 % | 20.0 % | 33.3 % | 40.0 % | 13.3% |
| Axin_466-480 **(4)** | 20.0 % | 13.3 % | 26.7 % | 6.7 % | 40.0 % | - |
| Axin_563-577 **(5)** | 6.7 % | 13.3 % | 26.7 % | 26.7 % | 13.3 % | 6.7 % |
| Axin_803-817 **(6)** | 6.7 % | 33.3 % | 20.0 % | 6.7 % | 26.7 % | - |
| **Clusterin_211-225 (7)** | - | 26.7 % | 20.0 % | 20 % | 33.3 % | 20.0 % |
| Clusterin_278-292 **(8)** | 26.7 % | 20.0 % | 26.7 % | 6.7 % | 13.3 % | - |
| Clusterin_338-352 **(9)** | 6.7 % | 26.7 % | 26.7 % | - | 20.0 % | 20.0 % |
| **αBcrystallin_8-22 (10)** | - | 20.0 % | 20.0 % | 26.7 % | 33.3 % | 26.7 % |
| αBcrystallin_137-151 **(11)** | 6.7 % | 13.3 % | 33.3 % | 20.0 % | 33.3 % | - |
| FZD9_41-55 **(12)** | 6.7 % | 6.7 % | 13.3 % | 20.0 % | 40.0 % | 6.7 % |
| DKK1_211-225 **(13)** | 6.7 % | 33.3 % | 20.0 % | 13.3 % | 26.7 % | - |
| HypF-N_12-23 **(14)** | - | 33.3 % | 8.3 % | 25.0 % | 25.0 % | 8.3 % |
| HypF-N_40-51 **(15)** | 25.0 % | 16.7 % | 8.3 % | 16.7 % | 25% | - |
| **SsoAcP_19-33 (16)** | - | 20.0 % | 6.7 % | 20.0 % | 33.3 % | 20.0 % |
| SsoAcP_57-71 **(17)** | 26.7 % | 13.3 % | 6.7 % | 6.7 % | 26.7 % | 6.7 % |
| mAcP_22-36 **(18)** | 20.0 % | 20.0 % | 6.7 % | 6.7 % | 26.7 % | 13.3 % |

* The residues are grouped into classes according to their nature and physical chemical properties: negatively charged (Asp, Glu), positively charged (Lys, Arg), hydrophilic (Ser, Thr, Asn, Gln, His), flexible (Gly, Pro), aliphatic (Ala, Cys, Val, Pro, Leu, Ile, Met), and aromatic (Phe, Tyr, Trp). The binders exerting an inhibitory activity are highlighted in red. In blue is highlighted the only non-inhibitory peptide displaying a similar composition as the inhibitors but a clustered distribution of the residues.

**Table S3.** Hydrophobic content of RGS L106R binders as calculated using the Roseman scale for hydrophobicity*.

| **Peptide** | **Hb** | **Hc** | **$Hb_N$** | **$Hc_N$** | **$Hb_N/Hc_N$** |
|---|---|---|---|---|---|
| Axin_338-352 **(1)** | + 8.39 | - 18.9 | + 0.559 | - 1.260 | 0.444 |
| Axin_368-382 **(2)** | + 10.68 | - 19.17 | + 0.712 | - 1.278 | 0.557 |
| Axin_436-450 **(3)** | + 11.17 | - 8.64 | + 0.745 | - 0.576 | 1.293 |
| Axin_466-480 **(4)** | + 9.05 | - 22.62 | + 0.603 | - 1.508 | 0.400 |
| Axin_563-577 **(5)** | + 4.94 | - 14.57 | + 0.329 | - 0.971 | 0.339 |
| Axin_803-817 **(6)** | + 7.28 | - 22.42 | + 0.485 | - 1.495 | 0.324 |
| **Clusterin_211-225 (7)** | **+ 13.42** | **- 17.14** | **+ 0.895** | **- 1.143** | **0.783** |
| Clusterin_278-292 **(8)** | + 3.12 | - 30.98 | + 0.208 | - 1.857 | 0.112 |
| Clusterin_338-352 **(9)** | + 9.67 | - 20.62 | + 0.645 | - 1.375 | 0.469 |
| **αBcrystallin_8-22 (10)** | **+ 14.72** | **- 14.97** | **+ 0.981** | **- 0.998** | **0.983** |
| αBcrystallin_137-151 **(11)** | + 7.23 | - 17.22 | + 0.482 | - 1.148 | 0.420 |
| FZD9_41-55 **(12)** | + 10.18 | - 9.77 | + 0.679 | - 0.651 | 1.043 |
| DKK1_211-225 **(13)** | + 5.41 | - 22.06 | + 0.361 | - 1.471 | 0.245 |
| HypF-N_12-23 **(14)** | + 6.69 | - 15.92 | + 0.558 | - 1.327 | 0.420 |
| HypF-N_40-51 **(15)** | + 4.42 | - 20.34 | + 0.368 | - 1.695 | 0.217 |
| **SsoAcP_19-33 (16)** | **+ 13.03** | **- 11.97** | **+ 0.869** | **- 0.798** | **1.089** |
| SsoAcP_57-71 **(17)** | + 8.23 | - 19.6 | + 0.549 | - 1.307 | 0.420 |
| mAcP_22-36 **(18)** | + 9.12 | - 21.3 | + 0.608 | - 1.420 | 0.428 |

* Hb and Hc are the hydrophilic and hydrophobic indexes prior to normalisation and are obtained as explained in the Materials and Methods section. $Hb_N$ and $Hc_N$ refer to the hydrophilic and hydrophobic indexes normalised by the total number of residues *N* present in each peptide (12 or 15 aa). The binders exerting an inhibitory activity are highlighted in red. In blue is highlighted the only non-inhibitory peptide displaying a similar composition as the inhibitors but a clustered distribution of the residues.

**Table S4.** Mean relative abundance and occurrence in inhibitory and non-inhibitory binders by class of amino acids*.

| | Negative | Positive | Hydro-philic | Flexible | Aliphatic | Aromatic |
|---|---|---|---|---|---|---|
| mean abundance in inhibitory binders | 0% | 20 % | 13 % | 23 % | 33 % | 23 % |
| mean abundance in non-inhibitory binders | 9 % | 20 % | 19 % | 16 % | 35% | 9 % |
| approx. mean enrichment | **- 9%** | 0 | **- 6%** | **+ 7 %** | -2 % | **+ 14%** |
| mean occurrence in inhibitory binders | **0** | 3 | **2** | **4** | 5.0 | **4** |
| mean occurrence in non-inhibitor binders | 1 | 3 | 3 | 2 | 5 | 1 |
| approx. residue enrichment | **- 1** | 0 | **- 1** | **+ 2** | 0 | **+ 3** |

* Enrichment (red) or depletion (blue) in the inhibitory binders composition are relative to that of non-inhibitory ones. All values are normalized for the number of residues of inhibitory (30 aa) or non-inhibitory binders (264 aa).

**Table S5.** Mean hydrophobic and hydrophilic content in inhibitory and non-inhibitory binders*.

| **Ensemble of peptides** | **Hb** | **Hc** | **$Hb_N$** | **$Hc_N$** | **$Hb_N/Hc_N$** |
|---|---|---|---|---|---|
| inhibitory binders | + 29.31 | - 26.94 | **+ 0.98** | - 0.90 | **1.09** |
| non-inhibitory binders | + 156.23 | - 328.21 | **+ 0.59** | - 1.24 | **0.48** |

* $Hb_N$ and $Hb_N/Hc_N$ values of the inhibitory binders (red) are highlighted as opposed to those of the non-inhibitory ones (blue). $Hb_N$ and $Hc_N$ were obtained by normalising Hb and Hc for the number of residues of inhibitory (30 aa) or non-inhibitory binders (264 aa).

**Table S6.** Relative abundance of amino acids in the chaperone-derived peptides designed to bear the molecular determinants of inhibitory activity*.

| **Peptide** | **Negative** | **Positive** | **Hydro-Philic** | **Flexible** | **Aliphatic** | **Aromatic** |
|---|---|---|---|---|---|---|
| αBcrystallin_45-61 **(19)** | - | 11.8 % | 17.6 % | 23.5 % | 35.3 % | 29.4 % |
| Clusterin_200-221 **(20)** | 4.5 % | 13.6 % | 22.7 % | 22.7 % | 31.8 % | 27.3 % |
| Clusterin_216-237 **(21)** | - | 9.1 % | 18.2 % | 22.7 % | 40.9 % | 22.7 % |

* The residues are grouped into classes according to their nature and physical chemical properties: negatively charged (Asp, Glu), positively charged (Lys, Arg), hydrophilic (Ser, Thr, Asn, Gln, His), flexible (Gly, Pro), aliphatic (Ala, Cys, Val, Pro, Leu, Ile, Met), and aromatic (Phe, Tyr, Trp).

**Table S7.** Hydrophobic content of the chaperone-derived peptides bearing the anti-aggregation molecular determinants binders as calculated using the Roseman scale for hydrophobicity*.

| **Peptide** | **Hb** | **Hc** | **$Hb_N$** | **$Hc_N$** | **$Hb_N/Hc_N$** |
|---|---|---|---|---|---|
| αBcrystallin_45-61 **(19)** | + 18.01 | - 11.62 | **+ 1.059** | - 0.684 | **1.548** |
| Clusterin_200-221 **(20)** | + 20.61 | - 14.87 | **+ 0.937** | - 0.676 | **1.386** |
| Clusterin_216-237 **(21)** | + 21.40 | - 17.94 | **+ 0.973** | - 0.815 | **1.194** |

* $Hb_N$ and $Hb_N/Hc_N$ (red) were kept near-constant to fit the molecular determinants. Hb and Hc are the hydrophilic and hydrophobic indexes prior to normalisation and are obtained as explained in the Materials and Methods section. $Hb_N$ and $Hc_N$ refer to the hydrophilic and hydrophobic indexes normalised by the total number of residues *N* present in each peptide (17 or 22 aa).

**Table S8.** Relative abundance of amino acids in the chaperone-derived peptides upon modulation of the molecular determinants of inhibitory activity*.

| **Peptide** | **Positive** | **Hydro-Philic** | **Flexible** | **Aliphatic** | **Aromatic** |
|---|---|---|---|---|---|
| **P**-SsoAcP_19-33 **(22)** | 20.0 % | 6.7 % | 20.0 % | 53.3 % | 20.0 % |
| **G**-αBcrystallin_8-22 **(23)** | 20.0 % | 20.0 % | 26.7 % | 6.7 % | 26.7 % |
| **W/T**-αBcrystallin_8-22 **(24)** | 20.0 % | 20.0 % | 26.7 % | 26.7 % | 33.3 % |
| **F/S**-αBcrystallin_8-22 **(25)** | 20.0 % | 20.0 % | 26.7 % | 26.7 % | 33.3 % |
| **W/T/G**-αBcrystallin_8-22 **(26)** | 20.0 % | 20.0 % | 26.7 % | - | 33.3 % |
| **L/S**-αBcrystallin_45-61 **(27)** | 11.8 % | 17.6 % | 23.5 % | 64.7 % | - |
| **F/S**-αBcrystallin_45-61 **(28)** | 11.8 % | 23.5 % | 23.5 % | 23.5 % | 41.2 % |

* The residues are grouped into classes according to their nature and physical chemical properties: negatively charged (Asp, Glu), positively charged (Lys, Arg), hydrophilic (Ser, Thr, Asn, Gln, His), flexible (Gly, Pro), aliphatic (Ala, Cys, Val, Pro, Leu, Ile, Met), and aromatic (Phe, Tyr, Trp).

**Table S9.** Hydrophobic content of the chaperone-derived upon modulation of the molecular determinants of inhibitory activity as calculated using the Roseman scale for hydrophobicity*.

| **Peptide** | **Hb** | **Hc** | **$Hb_N$** | **$Hc_N$** | **$Hb_N/Hc_N$** |
|---|---|---|---|---|---|
| **P**-SsoAcP_19-33 **(22)** | + 16.00 | - 11.97 | **+ 1.067** | - 0.798 | **1.337** |
| **G**-αBcrystallin_8-22 **(23)** | + 10.76 | - 14.97 | **+ 0.717** | - 0.998 | **0.718** |
| **W/T**-αBcrystallin_8-22 **(24)** | + 14.61 | - 14.85 | **+ 0.974** | - 0.990 | **0.984** |
| **F/S**-αBcrystallin_8-22 **(25)** | + 15.31 | - 15.57 | **+ 1.021** | - 1.038 | **0.984** |
| **W/T/G**-αBcrystallin_8-22 **(26)** | + 10.65 | - 14.85 | **+ 0.710** | - 0.990 | **0.717** |
| **L/S**-αBcrystallin_45-61 **(27)** | + 18.52 | - 11.62 | **+ 1.089** | - 0.684 | **1.592** |
| **F/S**-αBcrystallin_45-61 **(28)** | + 19.85 | - 12.86 | **+ 1.168** | - 0.756 | **1.545** |

* $Hb_N$ and $Hb_N/Hc_N$ (red) were varied within a relatively narrow range to modulate the molecular determinants. Hb and Hc are the hydrophilic and hydrophobic indexes prior to normalisation and are obtained as explained in the Materials and Methods section. $Hb_N$ and $Hc_N$ refer to the hydrophilic and hydrophobic indexes normalised by the total number of residues *N* present in each peptide (15 or 17 aa).

**Table S10.** Mean abundance of specific amino acids in RGS L106R binders and the inhibitors*.

| Residue | Relative abundance in the binders | Relative abundance in the inhibitors |
|---|---|---|
| Serine | 4.92 % | 6.67 % |
| Histidine | 4.92 % | 3.33 % |
| Glutamine | 3.03 % | 3.33 % |
| **Lysine** | **6.82 %** | **3.33 %** |
| **Arginine** | **13.64 %** | **16.67 %** |
| **Glycine** | **7.96 %** | **10.0 %** |
| **Proline** | **8.33 %** | **13.33 %** |
| **Valine** | **8.71 %** | **13.33 %** |
| **Leucine** | **7.58 %** | **3.33 %** |
| Isoleucine | 4.17 % | 3.33 % |
| Tyrosine | 3.03 % | 3.33 % |
| **Tryptophan** | **1.52 %** | **3.33 %** |
| **Phenylalanine** | **4.17 %** | **16.67 %** |

* In red are highlighted the residues that are enriched in the inhibitors, in blue those that are depleted and in green those that are slightly enriched despite being the strongest enhancers of the inhibitory activity. All the values are normalized for the total number of residues of inhibitors and non-inhibitory binders.

**Table S11**

**Peptide array 1**

The peptide array were screened against His-NUS-His, His-NUS-His-RGS WT or His-NUS-His-RGS L106R constructs. The protein bound on peptide spots was detected by anti-His HRP-antibody recognition and chemoluminescent ECL reagents.

| Position | | Residues | Sequence | Annotations |
|---|---|---|---|---|
| 1 | (A1) | | HHHHHHHH | His-tag |
| 2 | (A2) | 1-15 | MNIQEQGFPLDLGAS | Axin1 isoform1 |
| 3 | (A3) | 8-22 | FPLDLGASFTEDAPR | |
| 4 | (A4) | 16-30 | FTEDAPRPPVPGEEG | length: 15 |
| 5 | (A5) | 23-37 | PPVPGEEGELVSTDP | overlap: 8 |
| 6 | (A6) | 31-45 | ELVSTDPRPASYSFC | |
| 7 | (A7) | 38-52 | RPASYSFCSGKGVGI | |

| | | | |
|---|---|---|---|
| 8 | (A8) | 46-60 | SGKGVGIKGETSTAT |
| 9 | (A9) | 53-67 | KGETSTATPRRSDLD |
| 10 | (A10) | 61-75 | PRRSDLDLGYEPEGS |
| 11 | (A11) | 68-82 | LGYEPEGSASPTPPY |
| 12 | (A12) | 76-90 | ASPTPPYLKWAESLH |
| 13 | (A13) | 83-97 | LKWAESLHSLLDDQD |
| 14 | (A14) | 91-105 | SLLDDQDGISLFRTF |
| 15 | (A15) | 98-112 | GISLFRTFLKQEGCA |
| 16 | (A16) | 106-120 | LKQEGCADLLDFWFA |
| 17 | (A17) | 113-127 | DLLDFWFACTGFRKL |
| 18 | (A18) | 121-135 | CTGFRKLEPCDSNEE |
| 19 | (A19) | 128-142 | EPCDSNEEKRLKLAR |
| 20 | (A20) | 136-150 | KRLKLARAIYRKYIL |
| 21 | (A21) | 143-157 | AIYRKYILDNNGIVS |
| 22 | (A22) | 151-165 | DNNGIVSRQTKPATK |
| 23 | (A23) | 158-172 | RQTKPATKSFIKGCI |
| 24 | (A24) | 166-180 | SFIKGCIMKQLIDPA |
| 25 | (B1) | 173-187 | MKQLIDPAMFDQAQT |
| 26 | (B2) | 181-195 | MFDQAQTEIQATMEE |
| 27 | (B3) | 188-202 | EIQATMEENTYPSFL |
| 28 | (B4) | 196-210 | NTYPSFLKSDIYLEY |
| 29 | (B5) | 203-217 | KSDIYLEYTRTGSES |
| 30 | (B6) | 211-225 | TRTGSESPKVCSDQS |
| 31 | (B7) | 218-232 | PKVCSDQSSGSGTGK |
| 32 | (B8) | 226-240 | SGSGTGKGISGYLPT |
| 33 | (B9) | 233-247 | GISGYLPTLNEDEEW |

| | | | |
|---|---|---|---|
| 34 | (B10) | 241-255 | LNEDEEWKCDQDMDE |
| 35 | (B11) | 248-262 | KCDQDMDEDDGRDAA |
| 36 | (B12) | 256-270 | DDGRDAAPPGRLPQK |
| 37 | (B13) | 263-277 | PPGRLPQKLLLETAA |
| 38 | (B14) | 271-285 | LLLETAAPRVSSSRR |
| 39 | (B15) | 278-292 | PRVSSSRRYSEGREF |
| 40 | (B16) | 286-300 | YSEGREFRYGSWREP |
| 41 | (B17) | 293-307 | RYGSWREPVNPYYVN |
| 42 | (B18) | 301-315 | VNPYYVNAGYALAPA |
| 43 | (B19) | 308-322 | AGYALAPATSANDSE |
| 44 | (B20) | 316-330 | TSANDSEQQSLSSDA |
| 45 | (B21) | 323-337 | QQSLSSDADTLSLTD |
| 46 | (B22) | 331-345 | DTLSLTDSSVDGIPP |
| 47 | (B23) | 338-352 | SSVDGIPPYRIRKQH |
| 48 | (B24) | 346-360 | YRIRKQHRREMQESV |
| 49 | (C1) | 353-367 | RREMQESVQVNGRVP |
| 50 | (C2) | 361-375 | QVNGRVPLPHIPRTY |
| 51 | (C3) | 368-382 | LPHIPRTYRVPKEVR |
| 52 | (C4) | 376-390 | RVPKEVRVEPQKFAE |
| 53 | (C5) | 383-397 | VEPQKFAEELIHRLE |
| 54 | (C6) | 391-405 | ELIHRLEAVQRTREA |
| 55 | (C7) | 398-412 | AVQRTREAEEKLEER |
| 56 | (C8) | 406-420 | EEKLEERLKRVRMEE |
| 57 | (C9) | 413-427 | LKRVRMEEEGEDGDP |
| 58 | (C10) | 421-435 | EGEDGDPSSGPPGPC |
| 59 | (C11) | 428-442 | SSGPPGPCHKLPPAP |

| | | | |
|---|---|---|---|
| 60 | (C12) | 436-450 | HKLPPAPAWHHFPPR |
| 61 | (C13) | 443-457 | AWHHFPPRCVDMGCA |
| 62 | (C14) | 451-465 | CVDMGCAGLRDAHEE |
| 63 | (C15) | 458-472 | GLRDAHEENPESILD |
| 64 | (C16) | 466-480 | NPESILDEHVQRVLR |
| 65 | (C17) | 473-487 | EHVQRVLRTPGRQSP |
| 66 | (C18) | 481-495 | TPGRQSPGPGHRSPD |
| 67 | (C19) | 488-502 | GPGHRSPDSGHVAKM |
| 68 | (C20) | 496-510 | SGHVAKMPVALGGAA |
| 69 | (C21) | 503-517 | PVALGGAASGHGKHV |
| 70 | (C22) | 511-525 | SGHGKHVPKSGAKLD |
| 71 | (C23) | 518-532 | PKSGAKLDAAGLHHH |
| 72 | (C24) | 526-540 | AAGLHHHRHVHHHVH |
| 73 | (D1) | 533-547 | RHVHHHVHHSTARPK |
| 74 | (D2) | 541-555 | HSTARPKEQVEAEAT |
| 75 | (D3) | 548-562 | EQVEAEATRRAQSSF |
| 76 | (D4) | 556-570 | RRAQSSFAWGLEPHS |
| 77 | (D5) | 563-577 | AWGLEPHSHGARSRG |
| 78 | (D6) | 571-585 | HGARSRGYSESVGAA |
| 79 | (D7) | 578-592 | YSESVGAAPNASDGL |
| 80 | (D8) | 586-600 | PNASDGLAHSGKVGV |
| 81 | (D9) | 593-607 | AHSGKVGVACKRNAK |
| 82 | (D10) | 601-615 | ACKRNAKKAESGKSA |
| 83 | (D11) | 608-622 | KAESGKSASTEVPGA |
| 84 | (D12) | 616-630 | STEVPGASEDAEKNQ |
| 85 | (D13) | 623-637 | SEDAEKNQKIMQWII |

| | | | |
|---|---|---|---|
| 86 | (D14) | 631-645 | KIMQWIIEGEKEISR |
| 87 | (D15) | 638-652 | EGEKEISRHRRTGHG |
| 88 | (D16) | 646-660 | HRRTGHGSSGTRKPQ |
| 89 | (D17) | 653-667 | SSGTRKPQPHENSRP |
| 90 | (D18) | 661-675 | PHENSRPLSLEHPWA |
| 91 | (D19) | 668-682 | LSLEHPWAGPQLRTS |
| 92 | (D20) | 676-690 | GPQLRTSVQPSHLFI |
| 93 | (D21) | 683-697 | VQPSHLFIQDPTMPP |
| 94 | (D22) | 691-705 | QDPTMPPHPAPNPLT |
| 95 | (D23) | 698-712 | HPAPNPLTQLEEARR |
| 96 | (D24) | 706-720 | QLEEARRRLEEEEKR |
| 97 | (E1) | 713-727 | RLEEEEKRASRAPSK |
| 98 | (E2) | 721-735 | ASRAPSKQRYVQEVM |
| 99 | (E3) | 728-742 | QRYVQEVMRRGRACV |
| 100 | (E4) | 736-750 | RRGRACVRPACAPVL |
| 101 | (E5) | 743-757 | RPACAPVLHVVPAVS |
| 102 | (E6) | 751-765 | HVVPAVSDMELSETE |
| 103 | (E7) | 758-772 | DMELSETETRSQRKV |
| 104 | (E8) | 766-780 | TRSQRKVGGGSAQPC |
| 105 | (E9) | 773-787 | GGGSAQPCDSIVVAY |
| 106 | (E10) | 781-795 | DSIVVAYYFCGEPIP |
| 107 | (E11) | 788-802 | YFCGEPIPYRTLVRG |
| 108 | (E12) | 796-810 | YRTLVRGRAVTLGQF |
| 109 | (E13) | 803-817 | RAVTLGQFKELLTKK |
| 110 | (E14) | 811-825 | KELLTKKGSYRYYFK |
| 111 | (E15) | 818-832 | GSYRYYFKKVSDEFD |

| | | | | |
|---|---|---|---|---|
| 112 | (E16) | 826-840 | KVSDEFDCGVVFEEV | |
| 113 | (E17) | 833-847 | CGVVFEEVREDEAVL | |
| 114 | (E18) | 841-855 | REDEAVLPVFEEKII | |
| 115 | (E19) | 848-862 | PVFEEKIIGKVEKVD | |
| 116 | (E20) | 91-105 | SLLDDQDGISPFRTF | L101P |
| 117 | (E21) | 98-112 | GISLFMTFLKQEGCA | K103M |
| 118 | (E22) | 98-112 | GISLFRTFRKQEGCA | L106R |
| 119 | (E23) | 113-127 | DLLDFRRACTGFRKL | W118R/F119R |
| 120 | (E24) | 196-210 | NTYPSFLMSDIYLEY | R203M |
| 121 | (F1) | 608-622 | KAESSKSASTEVPGA | G612S |
| 122 | (F2) | 653-667 | SSGTRKPQLHENSRP | P661L |
| 123 | (F3) | 796-810 | YRTLVRGQAVTLGQF | R803Q |
| 124 | (F4) | 818-832 | GSYRYYKKKVSDEFD | F824K |
| 125 | (F5) | 818-832 | GSYRYYFKKVYDEFD | S828Y |
| 126 | (F6) | 833-847 | CGVVFEEVRKDEAVL | E842K |
| 127 | (F7) | 841-855 | REDEAVLLVFEEKII | P848L |
| 128 | (F8) | 848-862 | PVFEGKIIGKVEKVD | E852G |
| 129 | (F9) | 781-795 | DSIVVAYYRCGEPIP | F789R |
| 130 | (F10) | 788-802 | YFCGEPIPDRTLVRG | Y796D |
| 131 | (F11) | 803-817 | RAVTLGQRKELLTKK | F810R |
| 132 | (F12) | 818-832 | GSYRYYFAKVSDEFD | K825A |
| 133 | (F13) | 818-832 | GSYRYYFEKVSDEFD | K825E |
| 134 | (F14) | 818-832 | GSYRYYFKAVSDEFD | K826A |
| 135 | (F15) | 826-840 | KVSDEADCGVVFEEV | F831A |
| 136 | (F16) | 826-840 | KVSDEEDCGVVFEEV | F831E |
| 137 | (F17) | 826-840 | KVSDEFDCGVDFEEV | V836D |

| | | | | |
|---|---|---|---|---|
| 138 | (F18) | 833-847 | CGVVREEVREDEAVL | F837R |
| 139 | (F19) | 833-847 | CGVVFEEVREAEAVL | D843A |
| 140 | (F20) | 833-847 | CGVVFEEVREKEAVL | D843K |
| 141 | (F21) | 841-855 | REDEAVEPVFEEKII | L847E |
| 142 | (F22) | 1-15 | MMKTLLLFVGLLLTW | Clusterin |
| 143 | (F23) | 8-22 | FVGLLLTWESGQVLG | |
| 144 | (F24) | 16-30 | ESGQVLGDQTVSDNE | length: 15 |
| 145 | (G1) | 23-37 | DQTVSDNELQEMSNQ | overlap: 8 |
| 146 | (G2) | 31-45 | LQEMSNQGSKYVNKE | |
| 147 | (G3) | 38-52 | GSKYVNKEIQNAVNG | |
| 148 | (G4) | 46-60 | IQNAVNGVKQIKTLI | |
| 149 | (G5) | 53-67 | VKQIKTLIEKTNEER | |
| 150 | (G6) | 61-75 | EKTNEERKTLLSNLE | |
| 151 | (G7) | 68-82 | KTLLSNLEEAKKKKE | |
| 152 | (G8) | 76-90 | EAKKKKEDALNETRE | |
| 153 | (G9) | 83-97 | DALNETRESETKLKE | |
| 154 | (G10) | 91-105 | SETKLKELPGVCNET | |
| 155 | (G11) | 98-112 | LPGVCNETMMALWEE | |
| 156 | (G12) | 106-120 | MMALWEECKPCLKQT | |
| 157 | (G13) | 113-127 | CKPCLKQTCMKFYAR | |
| 158 | (G14) | 121-135 | CMKFYARVCRSGSGL | |
| 159 | (G15) | 128-142 | VCRSGSGLVGRQLEE | |
| 160 | (G16) | 136-150 | VGRQLEEFLNQSSPF | |
| 161 | (G17) | 143-157 | FLNQSSPFYFWMNGD | |
| 162 | (G18) | 151-165 | YFWMNGDRIDSLLEN | |
| 163 | (G19) | 158-172 | RIDSLLENDRQQTHM | |

| | | | |
|---|---|---|---|
| 164 | (G20) | 166-180 | DRQQTHMLDVMQDHF |
| 165 | (G21) | 173-187 | LDVMQDHFSRASSII |
| 166 | (G22) | 181-195 | SRASSIIDELFQDRF |
| 167 | (G23) | 188-202 | DELFQDRFFTREPQD |
| 168 | (G24) | 196-210 | FTREPQDTYHYLPFS |
| 169 | (H1) | 203-217 | TYHYLPFSLPHRRPH |
| 170 | (H2) | 211-225 | LPHRRPHFFFPKSRI |
| 171 | (H3) | 218-232 | FFFPKSRIVRSLMPF |
| 172 | (H4) | 226-240 | VRSLMPFSPYEPLNF |
| 173 | (H5) | 233-247 | SPYEPLNFHAMFQPF |
| 174 | (H6) | 241-255 | HAMFQPFLEMIHEAQ |
| 175 | (H7) | 248-262 | LEMIHEAQQAMDIHF |
| 176 | (H8) | 256-270 | QAMDIHFHSPAFQHP |
| 177 | (H9) | 263-277 | HSPAFQHPPTEFIRE |
| 178 | (H10) | 271-285 | PTEFIREGDDDRTVC |
| 179 | (H11) | 278-292 | GDDDRTVCREIRHNS |
| 180 | (H12) | 286-300 | REIRHNSTGCLRMKD |
| 181 | (H13) | 293-307 | TGCLRMKDQCDKCRE |
| 182 | (H14) | 301-315 | QCDKCREILSVDCST |
| 183 | (H15) | 308-322 | ILSVDCSTNNPSQAK |
| 184 | (H16) | 316-330 | NNPSQAKLRRELDES |
| 185 | (H17) | 323-337 | LRRELDESLQVAERL |
| 186 | (H18) | 331-345 | LQVAERLTRKYNELL |
| 187 | (H19) | 338-352 | TRKYNELLKSYQWKM |
| 188 | (H20) | 346-360 | KSYQWKMLNTSSLLE |
| 189 | (H21) | 353-367 | LNTSSLLEQLNEQFN |

| | | | | |
|---|---|---|---|---|
| 190 | (H22) | 361-375 | QLNEQFNWVSRLANL | |
| 191 | (H23) | 368-382 | WVSRLANLTQGEDQY | |
| 192 | (H24) | 376-390 | TQGEDQYYLRVTTVA | |
| 193 | (I1) | 383-397 | YLRVTTVASHTSDSD | |
| 194 | (I2) | 391-405 | SHTSDSDVPSGVTEV | |
| 195 | (I3) | 398-412 | VPSGVTEVVVKLFDS | |
| 196 | (I4) | 406-420 | VVKLFDSDPITVTVP | |
| 197 | (I5) | 413-427 | DPITVTVPVEVSRKN | |
| 198 | (I6) | 421-435 | VEVSRKNPKFMETVA | |
| 199 | (I7) | 428-442 | PKFMETVAEKALQEY | |
| 200 | (I8) | 436-449 | EKALQEYRKKHREE | |
| 201 | (I9) | 1-15 | MDIAIHHPWIRRPFF | αBcrystallin |
| 202 | (I10) | 8-22 | PWIRRPFFPFHSPSR | |
| 203 | (I11) | 16-30 | PFHSPSRLFDQFFGE | length: 15 |
| 204 | (I12) | 23-37 | LFDQFFGEHLLESDL | overlap: 8 |
| 205 | (I13) | 31-45 | HLLESDLFPTSTSLS | |
| 206 | (I14) | 38-52 | FPTSTSLSPFYLRPP | |
| 207 | (I15) | 46-60 | PFYLRPPSFLRAPSW | |
| 208 | (I16) | 53-67 | SFLRAPSWFDTGLSE | |
| 209 | (I17) | 61-75 | FDTGLSEMRLEKDRF | |
| 210 | (I18) | 68-82 | MRLEKDRFSVNLDVK | |
| 211 | (I19) | 76-90 | SVNLDVKHFSPEELK | |
| 212 | (I20) | 83-97 | HFSPEELKVKVLGDV | |
| 213 | (I21) | 91-105 | VKVLGDVIEVHGKHE | |
| 214 | (I22) | 98-112 | IEVHGKHEERQDEHG | |
| 215 | (I23) | 106-120 | ERQDEHGFISREFHR | |

| | | | | |
|---|---|---|---|---|
| 216 | (I24) | 113-127 | FISREFHRKYRIPAD | |
| 217 | (J1) | 121-136 | KYRIPADVDPLTITSS | |
| 218 | (J2) | 128-143 | VDPLTITSSLSSDGVL | |
| 219 | (J3) | 137-151 | LSSDGVLTVNGPRKQ | |
| 220 | (J4) | 144-159 | TVNGPRKQVSGPERTI | |
| 221 | (J5) | 152-167 | VSGPERTIPITREEKP | |
| 222 | (J6) | 160-175 | PITREEKPAVTAAPKK | |
| 223 | (J7) | 1-13 | DAEFRHDSGYEVH | Abeta42 |
| 224 | (J8) | 6-18 | HDSGYEVHHQKLV | |
| 225 | (J9) | 11-23 | EVHHQKLVFFAED | length: 13 |
| 226 | (J10) | 16-28 | KLVFFAEDVGSNK | overlap: 8 |
| 227 | (J11) | 21-33 | AEDVGSNKGAIIG | |
| 228 | (J12) | 26-38 | SNKGAIIGLMVGG | |
| 229 | (J13) | 31-42 | IIGLMVGGVVIA | |
| 230 | (J14) | 0-11 | GSAKNTSCGVQL | HypF-N |
| 231 | (J15) | 4-15 | NTSCGVQLNTSC | |
| 232 | (J16) | 8-19 | GVQLNTSCGVQL | length: 12 |
| 233 | (J17) | 4-15 | NTSCGVQLRIRG | overlap: 8 |
| 234 | (J18) | 8-19 | GVQLRIRGKVQG | |
| 235 | (J19) | 12-23 | RIRGKVQGVGFR | |
| 236 | (J20) | 16-27 | KVQGVGFRPFVW | |
| 237 | (J21) | 20-31 | VGFRPFVWQLAQ | |
| 238 | (J22) | 24-35 | PFVWQLAQQLNL | |
| 239 | (J23) | 28-39 | QLAQQLNLHGDV | |
| 240 | (J24) | 32-43 | QLNLHGDVCNDG | |
| 241 | (K1) | 36-47 | HGDVCNDGDGVE | |

| | | | | |
|---|---|---|---|---|
| 242 | (K2) | 40-51 | CNDGDGVEVRLR | |
| 243 | (K3) | 44-55 | DGVEVRLREDPE | |
| 244 | (K4) | 48-59 | VRLREDPETFLV | |
| 245 | (K5) | 52-63 | EDPETFLVQLYQ | |
| 246 | (K6) | 56-67 | TFLVQLYQHCPP | |
| 247 | (K7) | 60-71 | QLYQHCPPLARI | |
| 248 | (K8) | 64-75 | HCPPLARIDSVE | |
| 249 | (K9) | 68-79 | LARIDSVEREPF | |
| 250 | (K10) | 72-83 | DSVEREPFIWSQ | |
| 251 | (K11) | 76-87 | REPFIWSQLPTE | |
| 252 | (K12) | 80-91 | IWSQLPTEFTIR | |
| 253 | (K13) | 111-125 | PDHGYCQPISIPLCT | CRD Frizzled 1 |
| 254 | (K14) | 118-132 | PISIPLCTDIAYNQT | |
| 255 | (K15) | 126-140 | DIAYNQTIMPNLLGH | length: 15 |
| 256 | (K16) | 133-147 | IMPNLLGHTNQEDAG | overlap: 8 |
| 257 | (K17) | 141-155 | TNQEDAGLEVHQFYP | |
| 258 | (K18) | 148-160 | LEVHQFYPLVKVQ | |
| 259 | (K19) | 156-170 | LVKVQCSAELKFFLC | |
| 260 | (K20) | 161-175 | CSAELKFFLCSMYAP | |
| 261 | (K21) | 171-185 | SMYAPVCTVLEQALP | |
| 262 | (K22) | 176-192 | VCTVLEQALPPCRSLCE | |
| 263 | (K23) | 186-200 | PCRSLCERARQGCEA | |
| 264 | (K24) | 193-207 | RARQGCEALMNKFGF | |
| 265 | (L1) | 201-215 | LMNKFGFQWPDTLKC | |
| 266 | (L2) | 208-222 | QWPDTLKCEKFPVHG | |
| 267 | (L3) | 216-230 | EKFPVHGAGELCVGQ | |

| | | | | |
|---|---|---|---|---|
| 268 | (L4) | 23-37 | HSLFSCEPITLRMCQ | CRD Frizzled 3 |
| 269 | (L5) | 30-44 | PITLRMCQDLPYNTT | |
| 270 | (L6) | 38-52 | DLPYNTTFMPNLLNH | length: 15 |
| 271 | (L7) | 45-59 | FMPNLLNHYDQQTAA | overlap: 8 |
| 272 | (L8) | 53-67 | YDQQTAALAMEPFHP | |
| 273 | (L9) | 60-72 | LAMEPFHPMVNLD | |
| 274 | (L10) | 68-82 | MVNLDCSRDFRPFLC | |
| 275 | (L11) | 73-87 | CSRDFRPFLCALYAP | |
| 276 | (L12) | 83-97 | ALYAPICMEYGRVTL | |
| 277 | (L13) | 88-104 | ICMEYGRVTLPCRRLCQ | |
| 278 | (L14) | 98-112 | PCRRLCQRAYSECSK | |
| 279 | (L15) | 105-119 | RAYSECSKLMEMFGV | |
| 280 | (L16) | 113-127 | LMEMFGVPWPEDMEC | |
| 281 | (L17) | 120-136 | PWPEDMECSRFPDCDEP | |
| 282 | (L18) | 40-54 | EEERRCDPIRISMCQ | CRD Frizzled 4 |
| 283 | (L19) | 47-61 | PIRISMCQNLGYNVT | |
| 284 | (L20) | 55-69 | NLGYNVTKMPNLVGH | length: 15 |
| 285 | (L21) | 62-76 | KMPNLVGHELQTDAE | overlap: 8 |
| 286 | (L22) | 70-84 | ELQTDAELQLTTFTP | |
| 287 | (L23) | 77-89 | LQLTTFTPLIQYG | |
| 288 | (L24) | 85-99 | LIQYGCSSQLQFFLC | |
| 289 | (M1) | 90-104 | CSSQLQFFLCSVYVP | |
| 290 | (M2) | 100-114 | SVYVPMCTEKINIPI | |
| 291 | (M3) | 105-121 | MCTEKINIPIGPCGGMC | |
| 292 | (M4) | 115-129 | GPCGGMCLSVKRRCE | |
| 293 | (M5) | 122-136 | LSVKRRCEPVLKEFG | |

| | | | | |
|---|---|---|---|---|
| 294 | (M6) | 130-144 | PVLKEFGFAWPESLN | |
| 295 | (M7) | 137-151 | FAWPESLNCSKFPPQ | |
| 296 | (M8) | 145-161 | CSKFPPQNDHNHMCMEG | |
| 297 | (M9) | 28-42 | SKAPVCQEITVPMCR | CRD Frizzled 5 |
| 298 | (M10) | 35-49 | EITVPMCRGIGYNLT | |
| 299 | (M11) | 43-57 | GIGYNLTHMPNQFNH | length: 15 |
| 300 | (M12) | 50-64 | HMPNQFNHDTQDEAG | overlap: 8 |
| 301 | (M13) | 65-77 | LEVHQFWPLVEIQ | |
| 302 | (M14) | 73-87 | LVEIQCSPDLRFFLC | |
| 303 | (M15) | 78-92 | CSPDLRFFLCSMYTP | |
| 304 | (M16) | 88-102 | SMYTPICLPDYHKPL | |
| 305 | (M17) | 93-109 | ICLPDYHKPLPPCRSVC | |
| 306 | (M18) | 103-117 | PPCRSVCERAKAGCS | |
| 307 | (M19) | 110-124 | ERAKAGCSPLMRQYG | |
| 308 | (M20) | 118-132 | PLMRQYGFAWPERMS | |
| 309 | (M21) | 125-139 | FAWPERMSCDRLPVL | |
| 310 | (M22) | 133-150 | CDRLPVLGRDAEVLCMDY | |
| 311 | (M23) | 19-33 | HSLFTCEPITVPRCM | CRD Frizzled 6 |
| 312 | (M24) | 26-40 | PITVPRCMKMAYNMT | |
| 313 | (N1) | 34-48 | KMAYNMTFFPNLMGH | length: 15 |
| 314 | (N2) | 41-55 | FFPNLMGHYDQSIAA | overlap: 8 |
| 315 | (N3) | 49-63 | YDQSIAAVEMEHFLP | |
| 316 | (N4) | 56-68 | VEMEHFLPLANLE | |
| 317 | (N5) | 64-78 | LANLECSPNIETFLC | |
| 318 | (N6) | 69-83 | CSPNIETFLCKAFVP | |
| 319 | (N7) | 79-93 | KAFVPTCIEQIHVVP | |

| | | | | |
|---|---|---|---|---|
| 320 | (N8) | 84-100 | TCIEQIHVVPPCRKLCE | |
| 321 | (N9) | 94-108 | PCRKLCEKVYSDCKK | |
| 322 | (N10) | 101-115 | KVYSDCKKLIDTFGI | |
| 323 | (N11) | 109-123 | LIDTFGIRWPEELEC | |
| 324 | (N12) | 116-132 | RWPEELECDRLQYCDET | |
| 325 | (N13) | 34-58 | RGAAPCQAVEIPMCR | CRD Frizzled 9 |
| 326 | (N14) | 41-55 | AVEIPMCRGIGYNLT | |
| 327 | (N15) | 59-63 | GIGYNLTRMPNLLGH | length: 15 |
| 328 | (N16) | 56-70 | RMPNLLGHTSQGEAA | overlap: 8 |
| 329 | (N17) | 64-78 | TSQGEAAAELAEFAP | |
| 330 | (N18) | 71-83 | AELAEFAPLVQYG | |
| 331 | (N19) | 79-93 | LVQYGCHSHLRFFLC | |
| 332 | (N20) | 84-98 | CHSHLRFFLCSLYAP | |
| 333 | (N21) | 94-108 | SLYAPMCTDQVSTPI | |
| 334 | (N22) | 99-115 | MCTDQVSTPIPACRPMC | |
| 335 | (N23) | 109-123 | PACRPMCEQARLRCA | |
| 336 | (N24) | 116-130 | EQARLRCAPIMEQFN | |
| 337 | (O1) | 124-138 | PIMEQFNFGWPDSLD | |
| 338 | (O2) | 131-145 | FGWPDSLDCARLPTR | |
| 339 | (O3) | 139-155 | CARLPTRNDPHALCMEA | |
| 340 | (O4) | 85-99 | CAEDEECGTDEYCAS | CRD 1 DKK-1 |
| 341 | (O5) | 92-106 | GTDEYCASPTRGGDA | |
| 342 | (O6) | 100-114 | PTRGGDAGVQICLAC | length: 15 |
| 343 | (O7) | 107-121 | GVQICLACRKRRKRC | overlap: 8 |
| 344 | (O8) | 115-129 | RKRRKRCMRHAMCCP | |
| 345 | (O9) | 122-138 | MRHAMCCPGNYCKNGIC | |

| | | | | |
|---|---|---|---|---|
| 346 | (O10) | 189-203 | CLRSSDCASGLCCAR | CRD 2 DKK-1 |
| 347 | (O11) | 196-210 | ASGLCCARHFWSKIC | |
| 348 | (O12) | 204-218 | HFWSKICKPVLKEGQ | length: 15 |
| 349 | (O13) | 211-225 | KPVLKEGQVCTKHRR | overlap: 8 |
| 350 | (O14) | 219-233 | VCTKHRRKGSHGLEI | |
| 351 | (O15) | 226-240 | KGSHGLEIFQRCYCG | |
| 352 | (O16) | 234-248 | FQRCYCGEGLSCRIQ | |
| 353 | (O17) | 241-255 | EGLSCRIQKDHHQAS | |
| 354 | (O18) | 249-263 | KDHHQASNSSRLHTC | |
| 355 | (O19) | 147-161 | CIIDEDCGPSMYCQF | CRD 1 DKK-3 |
| 356 | (O20) | 154-168 | GPSMYCQFASFQYTC | |
| 357 | (O21) | 162-176 | ASFQYTCQPCRGQRM | length: 15 |
| 358 | (O22) | 169-183 | QPCRGQRMLCTRDSE | overlap: 8 |
| 359 | (O23) | 177-191 | LCTRDSECCGDQLCV | |
| 360 | (O24) | 184-195 | CCGDQLCVWGHC | |
| 361 | (P1) | 208-222 | CDNQRDCQPGLCCAF | CRD 2 DKK-3 |
| 362 | (P2) | 215-229 | QPGLCCAFQRGLLFP | |
| 363 | (P3) | 223-237 | QRGLLFPVCTPLPVE | length: 15 |
| 364 | (P4) | 230-244 | VCTPLPVEGELCHDP | overlap: 8 |
| 365 | (P5) | 238-252 | GELCHDPASRLLDLI | |
| 366 | (P6) | 245-259 | ASRLLDLITWELEPD | |
| 367 | (P7) | 253-267 | TWELEPDGALDRCPC | |
| 368 | (P8) | 260-274 | GALDRCPCASGLLCQ | |
| 369 | (P9) | 268-284 | ASGLLCQPHSHSLVYVC | |
| 370 | (P10) | 41-55 | CLSDTDCNTRKFCLQ | CRD 1 DKK-4 |
| 371 | (P11) | 48-62 | NTRKFCLQPRDEKPF | |

| | | | | |
|---|---|---|---|---|
| 372 | (P12) | 56-70 | PRDEKPFCATCRGLR | length: 15 |
| 373 | (P13) | 63-77 | CATCRGLRRRCQRDA | overlap: 8 |
| 374 | (P14) | 71-85 | RRCQRDAMCCPGTLC | |
| 375 | (P15) | 78-90 | MCCPGTLCVNDVC | |
| 376 | (P16) | 145-159 | CLRTFDCGPGLCCAR | CRD 2 DKK-4 |
| 377 | (P17) | 152-166 | GPGLCCARHFWTKIC | |
| 378 | (P18) | 160-174 | HFWTKICKPVLLEGQ | length: 15 |
| 379 | (P19) | 167-181 | KPVLLEGQVCSRRGH | overlap: 8 |
| 380 | (P20) | 175-189 | VCSRRGHKDTAQAPE | |
| 381 | (P21) | 182-196 | KDTAQAPEIFQRCDC | |
| 382 | (P22) | 190-204 | IFQRCDCGPGLLCRS | |
| 383 | (P23) | 197-211 | GPGLLCRSQLTSNRQ | |
| 384 | (P24) | 205-218 | QLTSNRQHARLRVC | |

**Peptide array 2**

The peptide array were screened against His-NUS-His, His-NUS-His-RGS WT or His-NUS-His-RGS L106R constructs. The protein bound on peptide spots was detected by anti-His HRP-antibody recognition and chemoluminescent ECL reagents.

| Position | | Protein | Residues | Sequence |
|---|---|---|---|---|
| 270 | (Q6) | mAcP | 0-14 | GSTAQSLKSVDYEVF |
| 271 | (Q7) | mAcP | 7-21 | KSVDYEVFGRVQGVS |
| 272 | (Q8) | mAcP | 15-29 | GRVQGVSFRMYTEDE |
| 273 | (Q9) | mAcP | 22-36 | FRMYTEDEARKIGVV |
| 274 | (Q10) | mAcP | 30-44 | ARKIGVVGWVKNTSK |
| 275 | (Q11) | mAcP | 37-51 | GWVKNTSKGTVTGQV |

| | | | | |
|---|---|---|---|---|
| 276 | (Q12) | mAcP | 45-59 | GTVTGQVQGPEDKVN |
| 277 | (Q13) | mAcP | 52-66 | QGPEDKVNSMKSWLS |
| 278 | (Q14) | mAcP | 60-74 | SMKSWLSKVGSPSSR |
| 279 | (Q15) | mAcP | 67-81 | KVGSPSSRIDRTNFSN |
| 280 | (Q16) | mAcP | 75-89 | IDRTNFSNEKTISKLE |
| 281 | (Q17) | mAcP | 82-98 | EKTISKLEYSNFSIRY |
| 282 | (Q18) | mAcP | 16-31 | RVQGVSFRMYTEDEAR |
| 283 | (Q19) | SsoAcP | -1-11 | GSMKKWSDTEVFE |
| 284 | (Q20) | SsoAcP | 12-26 | MLKRMYARVYGLVQG |
| 285 | (Q21) | SsoAcP | 19-33 | RVYGLVQGVGFRKFV |
| 286 | (Q22) | SsoAcP | 27-41 | VGFRKFVQIHAIRLG |
| 287 | (Q23) | SsoAcP | 34-48 | QIHAIRLGIKGYAKN |
| 288 | (Q24) | SsoAcP | 42-56 | IKGYAKNLPDGSVEV |
| 289 | (R1) | SsoAcP | 49-53 | LPDGSVEVVAEGYEE |
| 290 | (R2) | SsoAcP | 57-71 | VAEGYEEALSKLLER |
| 291 | (R3) | SsoAcP | 54-68 | ALSKLLERIKQGPPA |
| 292 | (R4) | SsoAcP | 72-86 | IKQGPPAAEVEKVDY |
| 293 | (R5) | SsoAcP | 69-83 | AEVEKVDYSFSEYKG |
| 294 | (R6) | SsoAcP | 87-101 | SFSEYKGEFEDFETY |
| 345 | (T9) | Fzd5 | 151-166 | NRSEATTAPPRPFPAK |
| 346 | (T10) | Fzd5 | 159-174 | PPRPFPAKPTLPGPPG |
| 347 | (T11) | Fzd5 | 167-182 | PTLPGPPGAPASGGEC |
| 348 | (T12) | Fzd5 | 175-190 | APASGGECPAGGPFVC |
| 349 | (T13) | Fzd5 | 183-198 | PAGGPFVCKCREPFVP |
| 350 | (T14) | Fzd5 | 191-206 | KCREPFVPILKESHPL |
| 351 | (T15) | Fzd5 | 199-214 | ILKESHPLYNKVRTGQ |

| | | | | |
|---|---|---|---|---|
| 352 | (T16) | Fzd5 | 207-222 | YNKVRTGQVPNCAVPC |
| 353 | (T17) | Fzd5 | 215-230 | VPNCAVPCYQPSFSAD |
| 354 | (T18) | Fzd5 | 223-238 | YQPSFSADERTFATFW |
| 355 | (T19) | Fzd5 | 292-307 | GHASVACSREHNHIHY |
| 356 | (T20) | Fzd5 | 300-315 | REHNHIHYETTGPALC |
| 357 | (T21) | Fzd5 | 380-395 | SVDGDPVAGICYVGNQ |
| 358 | (T22) | Fzd5 | 388-402 | GICYVGNQNLNSLRG |
| 359 | (T23) | Fzd5 | 471-486 | EQHYRESWEAALTCAC |
| 360 | (T24) | Fzd5 | 479-494 | EAALTCACPGHDTGQP |
| 361 | (U1) | Fzd5 | 487-502 | PGHDTGQPRAKPEYWV |
| 362 | (U2) | Fzd5 | 495-510 | RAKPEYWVLMLKYFMC |
| 363 | (U3) | Fzd5 | 503-518 | LMLKYFMCLVVGITSG |
| 364 | (U4) | Fzd5 | 511-526 | LVVGITSGVWIWSGKT |
| 365 | (U5) | Fzd5 | 519-534 | VWIWSGKTVESWRRFT |
| 366 | (U6) | Fzd5 | 527-542 | VESWRRFTSRCCCRPR |
| 367 | (U7) | Fzd5 | 535-551 | SRCCCRPRRGHKSGGAM |
| 368 | (U8) | Fzd5 | 543-559 | RGHKSGGAMAAGDYPEA |
| 369 | (U9) | Fzd5 | 552-568 | AAGDYPEASAALTGRTG |
| 370 | (U10) | Fzd5 | 560-576 | SAALTGRTGPPGPAATY |
| 371 | (U11) | Fzd5 | 569-585 | PPGPAATYHKQVSLSHV |
| 372 | (U12) | DKK1 | 32-46 | TLNSVLNSNAIKNLP |
| 373 | (U13) | DKK1 | 39-53 | SNAIKNLPPPLGGAA |
| 374 | (U14) | DKK1 | 47-61 | PPLGGAAGHPGSAVS |
| 375 | (U15) | DKK1 | 54-68 | GHPGSAVSAAPGILY |
| 376 | (U16) | DKK1 | 62-76 | AAPGILYPGGNKYQT |
| 377 | (U17) | DKK1 | 69-84 | PGGNKYQTIDNYQPYP |

| | | | | |
|---|---|---|---|---|
| 378 | (U18) | DKK1 | 139-153 | VSSDQNHFRGEIEET |
| 379 | (U19) | DKK1 | 146-160 | FRGEIEETITESFGN |
| 380 | (U20) | DKK1 | 154-168 | ITESFGNDHSTLDGY |
| 381 | (U21) | DKK1 | 161-175 | DHSTLDGYSRRTTLS |
| 382 | (U22) | DKK1 | 169-183 | SRRTTLSSKMYHTKG |
| 383 | (U23) | DKK1 | 176-188 | SKMYHTKGQEGSV |
| 384 | (U24) | His tag | | HHHHHH |
